# Retroreflective beam multiplexing enables high-throughput two-photon voltage imaging in vivo

**DOI:** 10.64898/2025.12.09.693349

**Authors:** Zheng Liu, Mengke Han, Yalan Chen, Nelson Patrick Downs, Jianglai Wu

## Abstract

Voltage imaging demands technical innovations to realize its full potential in neuroscience. Although two-photon microscopy is widely used in brain research, its imaging speed is insufficient for high-throughput voltage recordings. To overcome this limitation, we developed an all-optical beam multiplexing engine via retroreflection from two near-parallel mirrors. This approach integrates seamlessly with standard two-photon microscopes, enabling kilohertz-frame-rate imaging across a 16-fold expanded field of view while maintaining synaptic resolution. Operating at over 220 megapixels per second, it enables submillisecond imaging of dendritic propagation of action potentials and simultaneous voltage recording from more than 200 hippocampal neurons in head-fixed, behaving mice. This advancement unlocks new possibilities for large-scale, high-speed functional studies in neuroscience.

## Main

Monitoring the activity of large neuronal ensembles with high spatiotemporal resolution in vivo is crucial for unraveling the complex dynamics of neural circuits in behaving animals. Recent advances in voltage indicators have brought voltage imaging closer to becoming a transformative tool for neuroscience [1, 2]. Nevertheless, significant challenges remain, primarily due to the need for increasingly fast and sensitive imaging techniques capable of overcoming the highly scattering nature of brain tissues. Two-photon fluorescence microscopy (2PFM) addresses the intense tissue scattering via inherent optical sectioning and near-infrared excitation, enabling sharp, high-resolution imaging at substantial depths. Consequently, 2PFM has been routinely adopted for in vivo studies in neuroscience. For voltage imaging, however, achieving kilohertz (kHz) frame rate and enhancing throughput (effective pixel rate) is essential for studying activity across large neuronal populations. This requirement renders the widely adopted resonant-galvo scanner based 2PFM systems inadequate for large-scale voltage imaging, as the inertia-limited scanning mirrors cannot meet the necessary speed demand [3, 4].

Various laser scanning techniques have recently been developed to increase the speed of 2PFM for studying millisecond-scale neural dynamics in the living brain [5–10]. Among these techniques, the all-optical laser scanner based on free-space angular-chirp-enhanced delay (FACED) has achieved megahertz (MHz) line-scan rates, enabling full-frame two-photon voltage imaging of large neuronal populations in awake mice [10]. Nevertheless, the number of resolvable spots along the all-optical scanning axis is restricted to 80–100 because of instrumental limitations [11], and the obtainable image therefore is confined to a narrow strip with an equivalent number of lines. Although field of view tiling increases the number of lines in the image, it reduces the frame rate by a factor at least as large as the number of tiles. In addition, as each laser pulse is divided into 80–100 subpulses for all-optical scanning, efficient two-photon excitation necessitates low-repetition-rate (1–4 MHz) femtosecond lasers with microjoule pulse energies [9,10]. As a result, adapting the FACED all-optical laser scanner for labs equipped with conventional 2PFM remains challenge, as these systems typically use ∼80 MHz femtosecond lasers as excitation sources.

In this work, we introduce a FACED-enabled Retroreflective All-optical Multiplexing Engine (FRAME) to address these limitations. Seamlessly integrated with a commercial 2PFM platform, FRAME expands the field of view by 16-fold at kHz-frame-rate imaging, operating at the throughput limit imposed by fluorescence lifetime. We demonstrate the FRAME empowered high-throughput and high-resolution two-photon voltage imaging of neuronal populations in the primary visual cortex and hippocampus in awake mouse brains.

## RESULTS

The FACED device consists of two highly reflective, near-parallel mirrors with a tilt angle *α* (**Fig. 1a)**. A light ray entering at an incident angle of *nα* (where *n* is an integer) undergoes *n* internal reflections, then strikes one mirror at normal incidence and is retroreflected precisely along its original path [12]. Leveraging this retroreflective property, the previous FACED-2PFM platform transformed a femtosecond laser beam into 80–100 spatiotemporally evenly distributed beamlets for all-optical line scanning with MHz rates (**Fig. 1b**) — a design that inherently approaches the physical limit imposed by mirror size on the number of beamlets. In contrast, FRAME applies the same principle to realize spatiotemporal beam multiplexing, effectively avoiding the need to generate ever more beamlets and enabling higher-throughput imaging at kilohertz frame rate (**Fig. 1b**). In FRAME, the excitation source is an 80 MHz femtosecond laser (pulse-picked to 20 MHz). Using a diffractive beam splitter, the laser beam is split into a one-dimensional 16-beamlet array and coupled into the FACED device with a tunable tilt angle *α* and reconfigurable separation fixed at 468 mm (**Fig. 1a**, **Supplementary Fig. 1 and Methods**). The beamlet array has an inter-beam angle *α_s_* (∼2.4 mrad) at the device entrance point *O*. By tuning *α* to match *α_s_*, all beamlets undergo perfect retroreflection after multiple reflections, retracing their incident paths [12]. This process produces a sequence of sixteen subpulses (interpulse delay = 3.125 ns) evenly distributed within each period of the 20 MHz pulse train. The retroreflected beamlet array is delivered to a commercial 2PFM equipped with a 12 kHz resonant mirror for x-scanning and a galvanometer for y-scanning. In the objective focal plane, sixteen foci are formed and aligned along y-axis with 7.5 μm spacings, enabling multiplexed imaging with uniform optical resolution (x, 0.41 ± 0.01 µm; y, 0.41 ± 0.04 µm; z, 1.82 ± 0.13 µm; mean ± s.d. of 16 foci) and homogeneous illumination (< 5.2% power variations across all foci) (**Supplementary Figs. 2–3**). Each focus scans a 320 μm (max) × 7.5 μm subarea, collectively providing a field of view of 320 μm × 120 μm (**Fig. 1c**). When scanning the galvanometer unidirectionally at 805 Hz, conventional single-beam scanning produces 24-line images at 805 frames per second (fps). In contrast, FRAME expands the field of view by 16-fold without compromising the frame rate and image resolution, clearly resolving individual dendritic spines (**Fig. 1c**) and generating 384-line images—a 4 to 5-fold increase over FACED-2PFM (**Supplementary Table 1**).

**Fig. 1.**
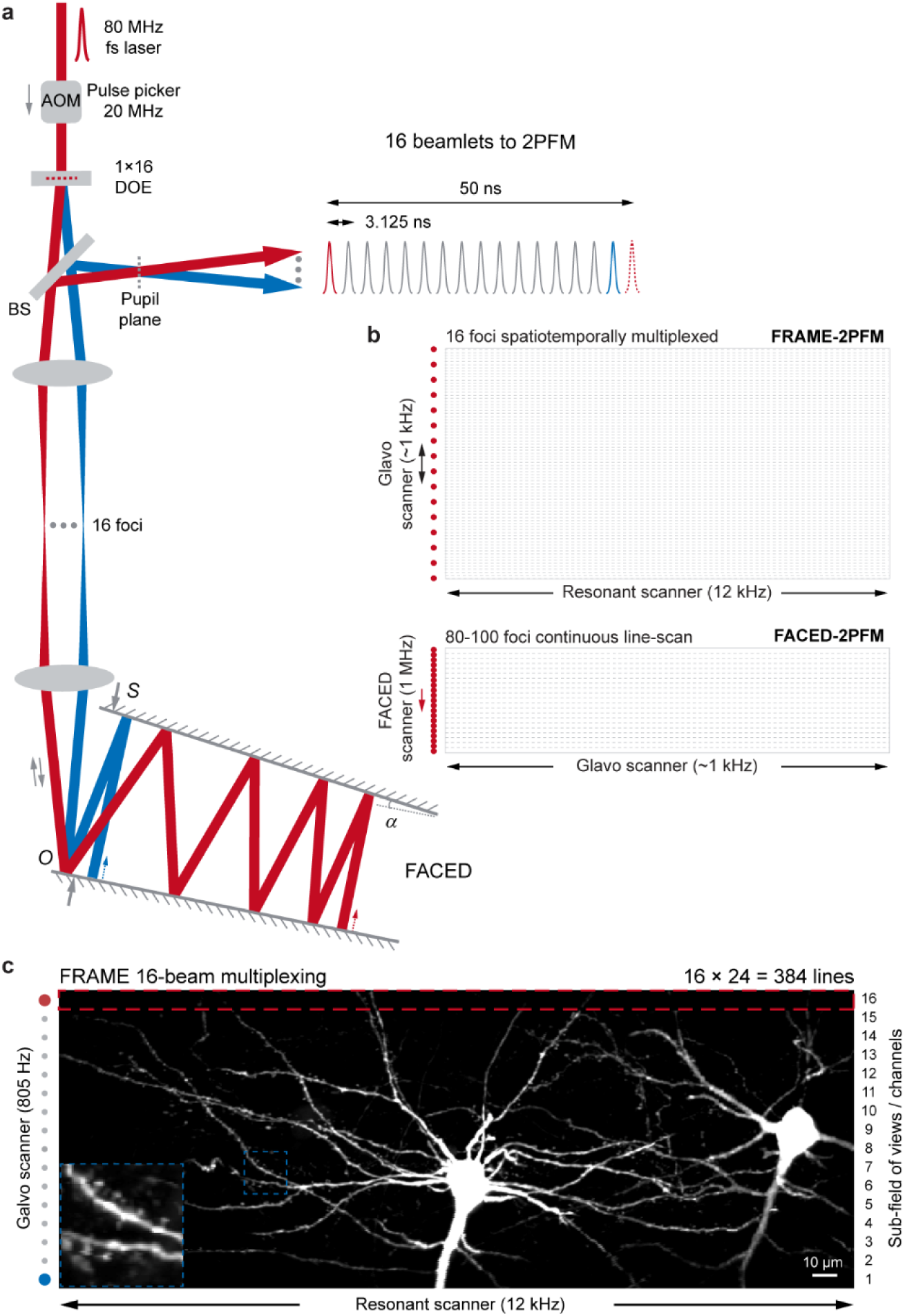
Working principle of FRAME-2PFM. a, Schematic of FRAME module. Retroreflection from the FACED mirror pair (*α*, tilt angle; *S*, mirror separation) temporally multiplexes the 16 beamlets split from a diffraction optical element (DOE); interpulse delay = 2S / c (c, light speed). A 4f optical system relays the pupil plane into the intermediate plane between the stacked galvo and resonant scanners of a standard 2PFM. AOM, acousto-optic modulator; BS, beam splitter. b, Beam multiplexing and scanning scheme in FACED-2PFM and FRAME-2PFM. c, Representative FRAME-2PFM image of a Thy1-eGFP mouse brain slice. The image is a maximum intensity projection of a 24-frame stack acquired at 1 μm intervals. The red dashed rectangle indicates the field of view achievable with conventional single-beam 2PFM. The region inside the dashed blue square is enlarged to show individual dendritic spines.

Using FRAME-2PFM, we first imaged the response of neurons in primary visual cortex (V1) to light flash in head-fixed awake mice (**Fig. 2a**). Neurons were sparsely expressing genetically encoded voltage indicator JEDI-2P-Kv [13]. The microscope enabled precise extraction of voltage signals from cell membrane-specific fluorescence (**Fig. 2b**). Typically, a population of 38 ± 6 neurons in V1 (mean ± s.d., n = 4 mice; **Supplementary Fig. 4**) were recorded in single fields of view. Imaging at 805 fps the microscope reliably captured both spiking and subthreshold activities (**Figs. 2c–d**), achieving an optical spike signal-to-noise ratio (SNR) of ∼6.0, with minimal fluorescence decay (< 10%) during continuous half-minute recordings (**Supplementary Figs. 5–6**). For light-responsive neurons, their spiking and subthreshold activities peaked after light stimulus (**Figs. 2e–f and Supplementary Fig. 7**), with a mean subthreshold response latency of 39.0 ± 2.4 ms (mean ± s.d., n = 202 cells from two mice; **Fig. 2g**), aligning with established electrophysiological observations [14,15]. We characterized the kinetics of optical spikes from this neuronal population. The measured full width at half-maximum was 2.93 ± 0.46 ms and the ΔF/F was 0.28 ± 0.05 (mean ± s.d., n = 195 neurons from two mice; **Supplementary Fig. 8**), consistent with previous characterizations [13].

**Fig. 2.**
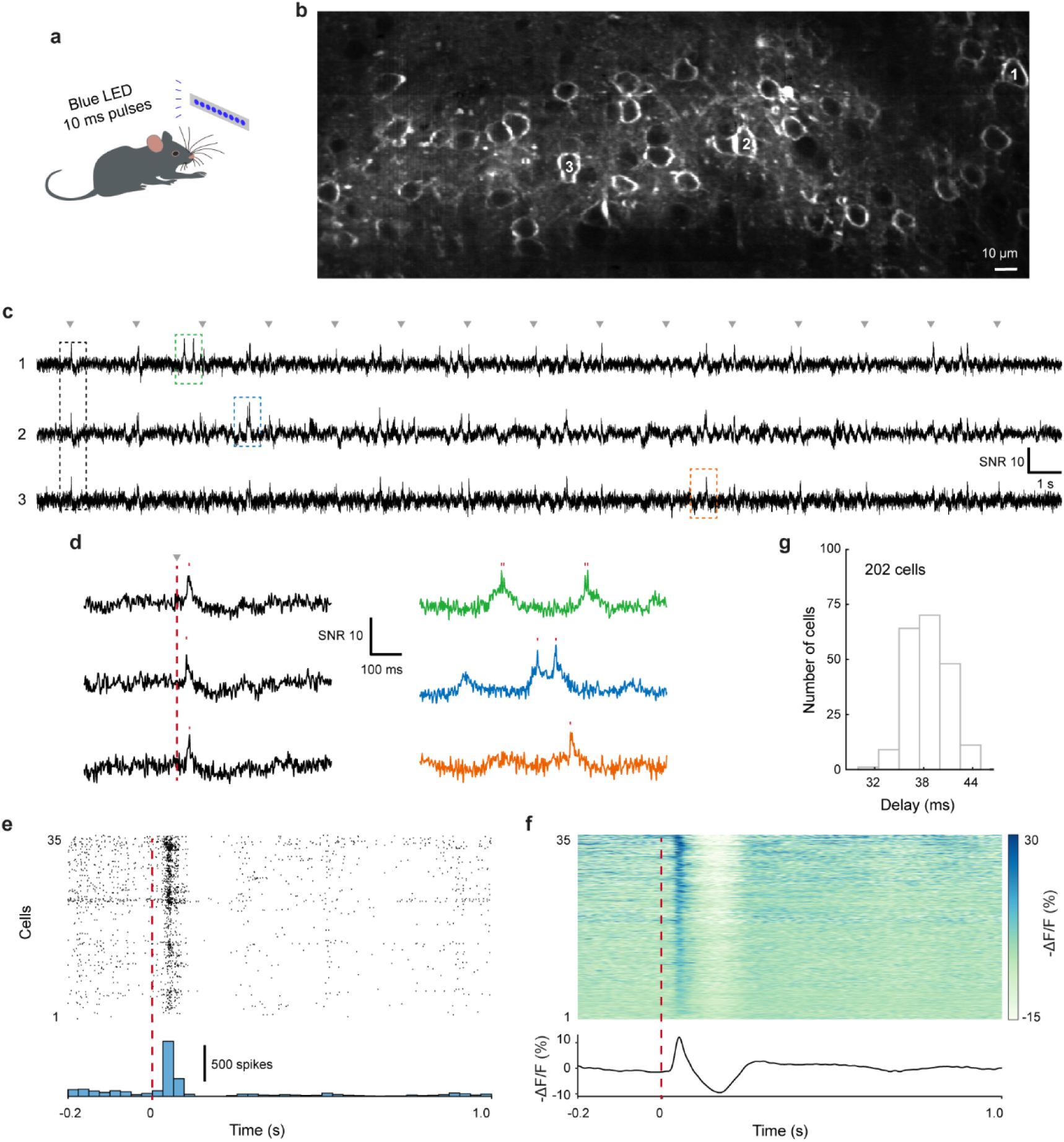
FRAME-2PFM in vivo voltage imaging in V1. **a**, Experimental paradigm. **b**, Representative image of V1 sparsely expressed with soma targeted JEDI-2P-Kv. **c**, Voltage traces from neurons labeled in (**b**); gray arrows indicate flash stimuli. **d,** Zoom-in view of color-boxed traces in (**c**); red ticks indicate spikes. **e–f**, Spiking (**e**) and subthreshold (**f**) responses of neurons in (**b**) (n = 135 stimuli). Top, spiking and subthreshold activities sorted by subthreshold response strength. Bottom, spike histogram and mean subthreshold strength. **g**, Subthreshold response latency of neuronal populations.

Dendritic voltage dynamics are essential for neuronal computation, impacting synaptic integration, plasticity, and the formation of complex output patterns. However, existing voltage imaging techniques have encountered difficulties in capturing rapid voltage signals in neuronal dendrites in vivo, mainly due to constraints in spatial resolution, speed, and sensitivity within opaque brain tissue. Leveraging the strength of two-photon imaging in highly scattering tissues, we next examined fast dendritic voltage dynamics in vivo at 1006 fps. FRAME-2PFM clearly resolved the dendritic membrane structures of a V1 neuron undergoing sustained firing (**Figs. 3a–b**), allowing somatic spike-triggered averaging to follow action potential as it propagated at ∼232 mm s^-1^ (**Figs. 3c–d and Methods**). This analysis revealed a ∼0.2 ms delay between somatic and dendritic spikes, underscoring FRAME-2PFM’s capability to resolve weak, submillisecond signals in fine neuronal structures in the living brain.

**Fig. 3.**
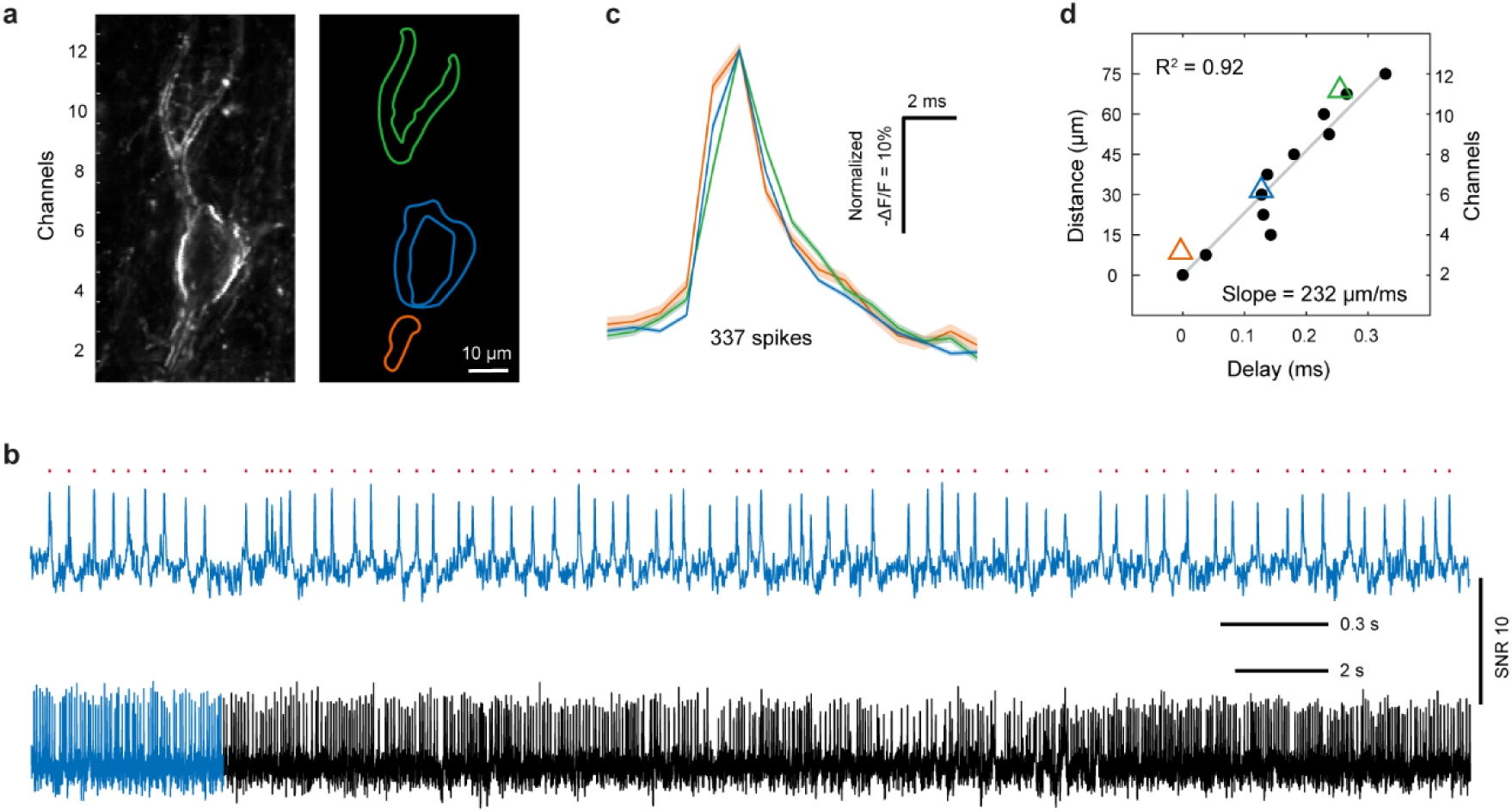
Imaging fast dendritic voltage dynamics using FRAME-2PFM. **a**, Image of a sustained-firing neuron (left) and masks of its soma / dendrites (right). The image shows a cropped region from a 96 μm × 120 μm field of view imaged at 1006 fps. **b**, Voltage trace from the soma (bottom), with a zoom-in view of region indicated in blue (top). **c**, Normalized optical spikes (mean ± s.e.m.) from the soma and dendrites labelled in (**a**). **d**, Action potential timing across different channels. Data points represent measured delays from the integrated signal in each channel; gray line shows their linear regression fit and triangles denote signal delays from the soma and dendrites labelled in (**a**).

Finally, we imaged the activities of neurons densely expressing JEDI-2P-Kv in hippocampus (CA1) of head fixed mice running on a treadmill (**Fig. 4a**). FRAME-2PFM enabled simultaneous recording from 203 ± 48 neurons (mean ± s.d., n = 4 mice; **Fig. 4b and Supplementary Fig. 9**), facilitating population-level analysis of network dynamics. In one animal, a subpopulation of the imaged neurons (32 / 150) exhibited highly synchronized spiking activity during locomotion (**Fig. 4c and Supplementary Fig. 10**). Pairwise cross-correlograms confirmed significantly stronger zero-lag synchronization during running compared to resting (**Fig. 4d and Methods**). Further spiking timing analysis revealed that neuronal firings were locked to theta-rhythm peaks (**Fig. 4e**), a phenomenon well-documented in electrophysiological observations [16]. Additionally, FRAME-2PFM enabled mapping of the preferred phase angles and locking strength of individual neurons (**Figs. 4f–g, Supplementary Fig. 11 and Methods**) with respect to their neighbors and anatomical landmarks. Population activity correlation and coherence analyses revealed distinct activity patterns between running and resting states, demonstrating state-dependent neural dynamics in CA1 (**Fig. 4h and Supplementary Fig. 12**). Moreover, single-cell spectral analysis showed diverse frequency responses across the neuronal population, with many neurons exhibiting peak activity in the gamma band (**Figs. 4i–j, Supplementary Figs. 13–14 and Methods**). Notably, imaging these high-frequency rhythms (> 40 Hz) at single-cell resolution remains challenging due to their low signal power [17].

**Fig. 4.**
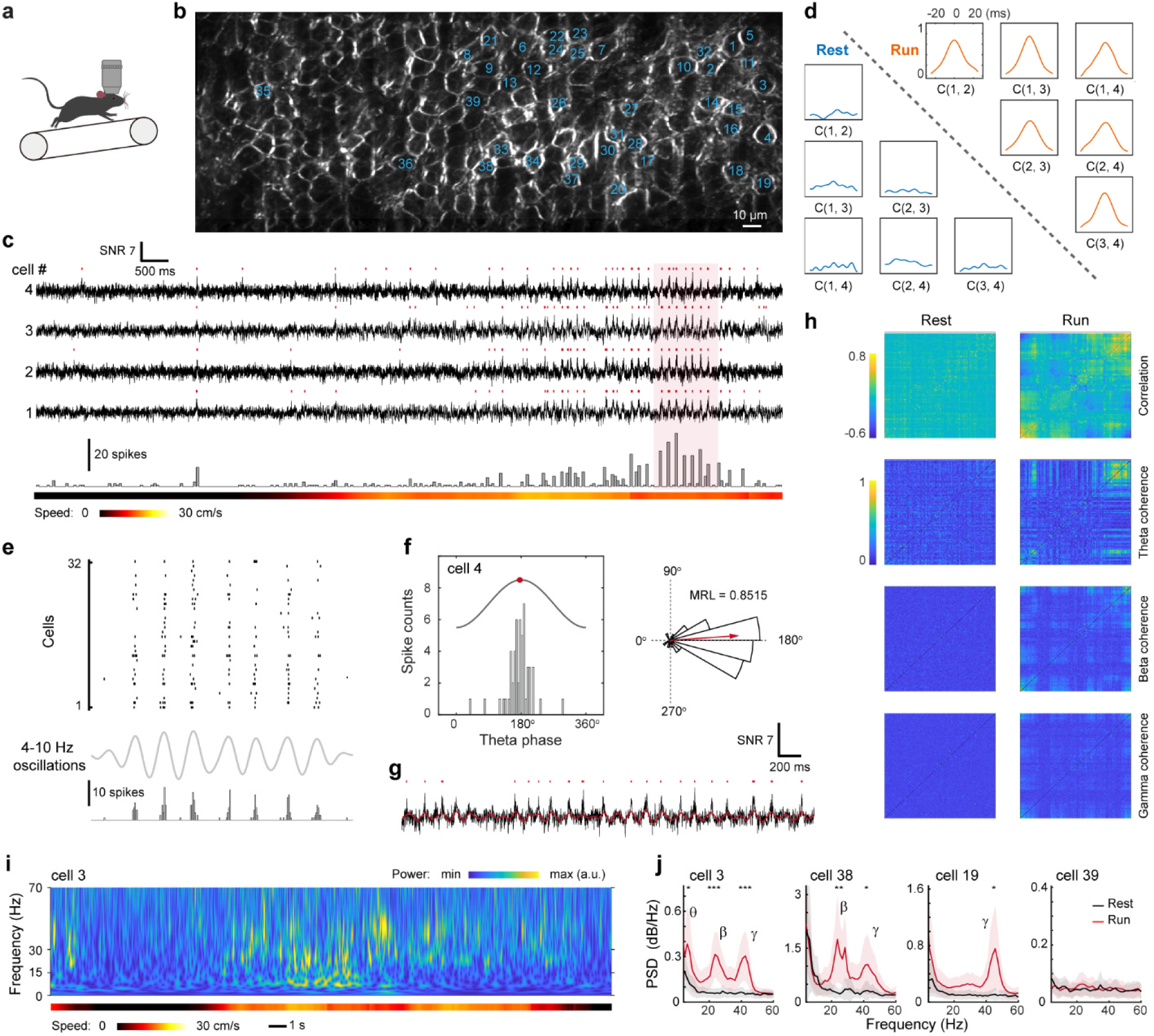
FRAME-2PFM in vivo voltage imaging in CA1. **a**, Experimental paradigm. **b**, Representative image of CA1 densely labeled with soma targeted JEDI-2P-Kv. **c**, Top, example traces from 4 of 32 neurons (numbered 1–32 in **b**) showing synchronized firing activity during running. Bottom, spike histogram from all running-modulated neurons along with running speed. **d**, Cross-correlograms derived from the traces in (**c)** during running and resting states. **e,** Spike trains of the 32 neurons (top), theta-band filtered trace (middle), and summed spiking activity across the population (bottom) during the running epoch shaded in (**c**). **f**, Left, spike distribution within a theta cycle for an example neuron; red dot marks the median phase (176.2°). Right, polar plot of spike phases (18° bin size); mean resultant length (MRL, range 0–1) indicates the phase locking strength. **g**, Expanded voltage trace and corresponding theta-oscillation profile (red line) from neuron in (**f**) during running. **h**, Correlation map (top) and coherence map across frequency bands (bottom) for all 150 neurons imaged during running and resting states. Neuronal order was determined by hierarchical clustering of the running-state correlation matrix and applied to all maps. **i**, Frequency spectrogram along with running speed of an example neuron. **j,** Power spectra of example neurons during running and resting states (mean ± s.d., n = 6 trials); PSD, power spectral density; two-tailed non-parametric Wilcoxon rank-sum test: *p < 0.05, **p < 0.01, and ***p < 0.001.

## DISCUSSION

In summary, FRAME achieved a line scan rate of 384 kHz and maximum throughput of 320 megapixels per second (pixel dwell time = 3.125 ns) limited by fluorescence lifetime (**Supplementary Fig. 15**), enabling high-throughput voltage imaging on a conventional 2PFM. All in vivo imaging was performed using safe laser power below 200 mW (**Supplementary Table 2**). For sparse labelling as in V1, using a digital micromirror device to mask out the area without neurons would increase the efficiency of power usage, akin to the adaptive illumination two-photon imaging technique [8]. As in FACED-2PFM [9, 10], the high-throughput operation in FRAME ensures each pixel is excited by a single laser pulse. This single-pulse excitation enhances photon yield from individual fluorophores by providing extended recovery periods, facilitating their transition from damage-susceptible dark states back to ground state [18].

Previously, the FACED all-optical scanner for two-photon voltage imaging creates a continuous line scan comprising 80 pixels, a count limited by the custom mirror size, laser pulse energy, and fluorescence lifetimes [9, 11]. FACED 2.0 increases this pixel count to 100 per line with the same hardware by halving the mirror separation from 30 cm to 15 cm [10]. Consequently, the interpulse delay decreases from 2 ns to 1 ns, which substantially increases the fluorescence crosstalk among adjacent pixels (**Supplementary Fig. 15**). In addition, beamlets along the line scan originate from virtual light sources with equal fan angles but different path lengths, causing their beam sizes to increase monotonically from the first to the last beamlet at the pupil plane [11]. Ensuring a uniform PSF across the line necessitates vignetting the largest beamlet to the size of the smallest, which unavoidably results in optical power loss. FRAME circumvents these limitations by exploiting the retroreflective properties of a FACED device for spatiotemporal beam multiplexing. Rather than densely packing beamlets into a continuous line, it generates 16 discrete, multiplexed beamlets (**Fig. 1b**). This operation directly multiplies the resonant scanner’s 24-kHz base line scan rate and thereby significantly increases the effective number of pixels at high-speed imaging, enhancing both image resolution and sensitivity for demanding applications like voltage imaging.

FRAME is a passive module built with off-the-shelf components (**Supplementary Table 3**). It effectively folds an extended optical path of approximately 14 meters, spanning from the first to the sixteenth beamlet, into a compact footprint of around 300 mm × 500 mm. In contrast to conventional time-multiplexing methods, which require substantially more optical table space for comparable path lengths, this design offers a significantly more compact and practical solution. Moreover, FRAME can be added onto a standard resonant-galvo scanner based 2PFM equipped with the widely used ∼80 MHz femtosecond lasers. Notably, spatiotemporal multiplexing in FRAME does not sacrifice the field of view along resonant scanning axis, differing from techniques like the scan multiplier unit and optical gear box [5, 6], where line scan from a resonant or polygon mirror scanner is divided into multiple shorter line scans to increase scan rate. In addition, replicating the sixteen multiplexed beamlets to simultaneously image two neighboring fields of view with multichannel photomultiplier tube detection [7], we could double the throughput. As throughput scales, however, boosting laser power becomes critical to maintain the SNR. Interpulse delay in FRAME is optimized to ensure high throughput while minimizing signal crosstalk. We note that the residual crosstalk diminishes the SNR by introducing additional photon noise, especially in densely labeled samples (**Supplementary Fig.5**). Increasing interpulse delay reduces signal crosstalk. Counterintuitively, this delay extension does not necessitate greater mirror separation—a modification that would otherwise enlarge the device footprint. For example, in current FRAME module, masking the odd or even numbered beamlets realizes 8-beamlet multiplexing and the interpulse delay is extended to 6.25 ns. This operation would halve the line scan rate and throughput but substantially lower signal crosstalk.

Compared to one-photon voltage imaging, the two-photon approach requires higher laser power, imposing stricter optical constraints to ensure tissue safety [19]. Consequently, when combined with sparse labeling or targeted illumination, one-photon voltage imaging captures larger neuronal populations in vivo using advanced specialized confocal microscopes [20, 21]. However, we expect FRAME-2PFM to maintain its niche in studying densely labeled specimens, as demonstrated in hippocampal imaging. In these cases, one-photon voltage imaging would face significant challenges to extract the signal from strong fluorescence background. Future advancements in optical instrumentation, along with better voltage indicators and computational algorithms are expected to unlock the full potential of two-photon voltage imaging.

## METHODS

### Animals

All experimental procedures and protocols on mice were approved by the Institutional Animal Care and Use Committee of the Chinese Institute for Brain Research, Beijing (CIBR) and complied with the governmental regulations of China.

### Microscope design

The schematic of FRAME-2PFM is shown in **Supplementary Fig. 1a**. The original 80 MHz repetition rate of the two-photon excitation laser (Insight X3+, Spectra Physics;

∼120-fs pulse width) was reduced to 20 MHz through pulse picking using an acousto-optic modulator (AOM; MT350-A0.12-800, AA Opto-electronic). The laser beam was first focused into the AOM using an achromatic doublet (AC254-050-B-ML, Thorlabs), then expanded to ∼3.6 mm diameter with a second achromatic doublet (AC254-150-B-ML, Thorlabs). Synchronization between the AOM and laser system is detailed in the *Synchronization Control* section. Following pulse picking, the laser beam was transformed into a one-dimensional 16-beamlet array using a diffraction optical element (DOE; MS-770-L-Y-A, HOLO/OR; separation angle = 0.22°). This array was then coupled into the FACED device through a 4f relay system formed by two achromatic doublets (AC508-250-B-ML and AC508-400-B-ML, Thorlabs) to produce an inter-beam angle of 2.4 mrad. The FACED device consisted of a near-parallel mirror pair (reflectivity > 99.9% at 900–1050 nm, fused silica substrate, 250 mm long and 20 mm wide, Layertec) with separation and tilt angle configured to ∼468 mm and ∼2.4 mrad, enabling perfect retroreflection of all beamlets after multiple reflections to generate 16 spatiotemporally multiplexed beamlets. A polarization beam splitter (CCM1-PBS252, Thorlabs) in combination with a half-wave plate (AHWP05M-980, Thorlabs) and quarter-wave plate (AQWP10M-980, Thorlabs) were used to separate the incident and retroreflected beamlet array. The FRAME module demonstrated a power throughput of ∼68%, calculated as the ratio between power measured at pupil plane to the power measured immediately after AOM (**Fig. 1a**).

The pupil plane of the FRAME module was relayed to the intermediate plane between the stacked x- and y- scanners (LSK-GR12, Thorlabs) in the commercial 2PFM (Bergamo II, Thorlabs) using an additional 4f system composed of two achromatic doublets (AC508-300-B-ML and AC508-250-B-ML, Thorlabs). Following the microscope’s built-in scan and tube lenses, the output beam (∼12 mm in diameter) underfilled the back aperture of the 25× 1.05 NA water-dipping objective (XLPLN25XWMP2, Olympus). In the focal plane, sixteen foci were generated with 7.5 μm y-axis spacing, covering a total field of view of ∼320 µm (x-axis, limited by scan range of the 12 kHz resonant scanner) × 120 µm (y-axis). Fluorescence signals were detected using an amplified non-cooled GaAsP PMT (PMT3100R, Thorlabs) and sampled at 2.56 GHz with a high-speed digitizer (High-Speed vDAQ, MBF Bioscience). Real-time data can be streamed to the host computer continuously for more than two minutes without exceeding the data transfer bandwidth. All microscope control, data acquisition, and image demultiplexing were performed using ScanImage (MBF Bioscience) running on MATLAB 2022b (MathWorks). The implementation of FRAME boosted the line scan rate from 24 kHz to 384 kHz. Using unidirectional Y-galvo scanning at 805 Hz and 1006 Hz, the microscope achieved frame rates of 805 fps (384 lines/frame) and 1006 fps (288 lines/frame), sustaining a continuous acquisition rate of 228–243 megapixels per second with 787 pixels per line (one pixel per subpulse).

### Synchronization control

The synchronization control scheme is illustrated in **Supplementary Fig. 1b**. The 80 MHz analog output from the laser was first converted to a square wave via a voltage comparator and frequency divided to yield a 10 MHz reference clock. This reference synchronized a waveform generator (AFG31102, Tektronix) to produce 20 MHz square waves (28% duty cycle; pulse width 14 ns) phase- locked to the laser output. These signals served as both the AOM gate for pulse picking and the external clock for the high-speed digitizer. FRAME-2PFM leveraged the virtual channel multiplexing function in ScanImage to demultiplex the signals excited from the spatiotemporally multiplexed 16-beamlets and to produce 16 sub-channel images for each 2D scan. These sub-channel images were then stitched together to form a final image using a custom MATLAB script.

### Point spread function characterization

The PSF of FRAME-2PFM was measured using Ø 200 nm fluorescent beads (Invitrogen, F8811). A 0.35 μL aliquot of the bead suspension was diluted in 1 mL of molten 0.5% agarose (Meilun, MB2534) and cast into a thin film in a 35 mm petri dish. The solidified gel was dried on a 40°C hotplate to immobilize the beads prior to imaging. For each beamlet, 3D image stacks of isolated beads were acquired within its sub-field of view at a voxel size of 58 nm × 58 nm × 221 nm. The intensity profiles through the bead centers were fitted with Gaussian functions along each axis. The PSF, defined as the full-width-at-half-maximum (FWHM) of the fitted profiles, is reported as the average from multiple beads.

### Mouse preparation for in vivo imaging

Male and female C57BL/6J mice (Jackson Laboratory, stock #000664) aged 6–7 weeks were used in this study. Mice were group-housed (1–5 per cage) under a reversed light cycle before surgery and singly or pair-housed afterward. For all surgical procedures, mice were anesthetized with isoflurane (3% induction, 1–1.5% maintenance) and head-fixed in a stereotaxic apparatus. Viral injections and subsequent cranial window (V1) or imaging cannula (CA1) implantations were performed according to established protocols [9, 22, 23]. The AAV2/9-hSyn-JEDI-2P-Kv vector (titer: 3.51 × 10^12^ GC/mL; Taitool Bioscience, China) was used to label both V1 and CA1 neurons.

*V1-specific preparation.* A 3.8-mm-diameter craniotomy was performed over left V1 with dura left intact. A glass capillary was beveled at 45° to create a 15–20 μm opening, back-filled with mineral oil, and fitted with a plunger controlled by a microinjection pump (R-480, RWD Life Science). This apparatus was used to inject (0.4 nL / sec) the viral solution into the brain to a depth of ∼ 200–400 μm below the pia. Virus was injected at three sites (75 nL per site) with 0.3–0.5 mm spacing between sites. After viral injection, a glass window (3 mm diameter, 0.15 mm thickness) was embedded in the craniotomy and secured with Vetbond (3M, USA). Finally, a custom titanium baseplate was cemented to the skull with dental acrylic to permit head-fixation.

*CA1-specific preparation*. A 3-mm-diameter craniotomy was performed above left dorsal CA1. Viral solution was injected at two sites (100 nL per site) using similar method described for V1. The overlying cortical tissue was aspirated using blunt 27- and 28-gauge needles connected to a vacuum under continuous irrigation with 4°C sterile saline. Once the corpus callosum was visible, large superficial fibers were removed with a blunt 30-gauge needle, leaving the thin underlying fiber layer intact. A custom imaging cannula (stainless steel ring: 3 mm outer diameter, 0.1 mm wall thickness, 1.2 mm height) with a bonded glass base (3 mm diameter, 0.15 mm thickness) was slowly and vertically advanced until the glass contacted the fiber layer. The cannula was then secured with Vetbond (3M, USA), and a custom titanium headplate was attached with dental acrylic for head-fixation.

Following surgery, animals were removed from the stereotaxic frame and placed on a 37°C heating pad to maintain body temperature during recovery. After a minimum two-week recovery period, mice were habituated to head-fixation prior to in vivo imaging.

### Visual stimulation in head-fixed awake mice

Visual stimuli were delivered via a blue LED strip (20 cm × 0.5 cm) positioned 15 cm from the mouse’s right eye at a ∼40° angle relative to its long axis. Head-fixed mice were restrained in an acrylic tube to limit movement. For each trial, imaging began 1 second before stimulus onset. Each stimulus consisted of a 10 ms light flash followed by a 2-second inter-stimulus interval, repeated 15 times per trial. Each experimental session comprised 9–10 trials, with trials lasting ∼30 seconds (25, 000 frames).

### Hippocampal imaging on a treadmill

Before imaging, mice were acclimated to the treadmill with one 2-hour training session per day for one week. During imaging, mice were head-fixed and positioned above a 20-cm-long custom treadmill with an Oxford fabric belt that permitted voluntary locomotion. A rotary encoder tracked belt movement to obtain the animal’s instantaneous position, and locomotion speed was extracted offline from this data. Each experimental session comprised 6-8 trials, with trials lasting ∼30 seconds (25, 000 frames).

### Signal crosstalk correction

Crosstalk coefficients for four adjacent channels were derived from the fluorescence decay curve (**Supplementary Fig. 15**). Each coefficient was defined as the ratio of the intensity integral for a given channel to that of the first channel, yielding values of 1, 0.37, 0.11, and 0.02. We subsequently applied a pixel-wise correction algorithm that processed each frame by sequentially analyzing groups of four pixels along y-axis. For each group, the algorithm utilized these coefficients to subtract the estimated crosstalk contribution from channels 2–4 and reassign it to the first channel.

### Analysis of activity data

Motion correction was performed with the NoRMCorre algorithm [24]. Periods with uncorrectable motion artifacts were discarded. We manually selected region of interests (ROIs) from the averaged images of the registered image sequence. Note that in hippocampal recordings, some ROIs unavoidably contained signals from multiple (∼1–3) neighboring neurons due to dense packing. The mean fluorescence intensity within the ROIs was used to calculate the functional traces. All analyses were conducted in MATLAB 2024a utilizing a suite of functions as specified below.

*Spike detection.* First, the raw fluorescence trace was corrected for photobleaching by subtracting its baseline, which was obtained by applying a moving median filter with a 500 ms window. Then, the detrended fluorescence trace was filtered using a 250 Hz 5th-order Butterworth low-pass filter to isolate suprathreshold activity. All optically recorded voltage traces presented in the manuscript exhibit suprathreshold activity. Spike detection was performed by identifying peaks with a minimum inter-spike interval of 3 ms. For SNR calculation, noise was defined as the standard deviation of a 20 Hz-high-pass-filtered version of the suprathreshold activity across the entire imaging session. Signal was defined as the peak value of a spike, with an initial SNR threshold set at four times the noise level. This threshold was manually adjusted as needed, and all traces were visually inspected to confirm the accuracy of spike detection. For SNR statistics, the maximum SNR of a trial was used as the corresponding SNR for each neuron. The relative fluorescence change (ΔF/F) for each neuron was calculated as the mean fractional fluorescence decrease at the peak of all detected spikes, with F taken as the mean of the baseline fluorescence.

*Action potential timing.* Propagation delay of action potential was calculated using the SNAPT algorithm [25]. Briefly, somatic spikes were identified and used as temporal references to generate a spike-triggered averaged movie (21 frames), which were then mean-filtered (radius = 2) using ImageJ. Spike timing at each pixel (with membrane structures) was defined as the time when the fluorescence trace first crossed a threshold of 60% of the maximum fluorescence change. These spike times were then averaged for each channel. Propagation delay across channels, along with propagation delay determined from the three ROIs in **Fig. 3a**, were displayed in **Fig. 3d**.

*Subthreshold oscillations extraction.* Subthreshold activity was obtained by applying a 50 Hz 5th-order Butterworth low-pass filter to the detrended fluorescence trace. To depict the time-frequency characteristics of the subthreshold activity, a Morlet-wavelet transform was applied to the raw fluorescence trace, yielding a frequency-resolved spectrogram (**Fig. 4i and Supplementary Fig. 13**). Single-trial fluorescence traces illustrating membrane potential oscillations within different bands were extracted by applying a 3rd-order Butterworth bandpass filter to the raw fluorescence traces. Power spectral density (PSD) was estimated by applying the “pwelch” function (500 ms Hamming window, 50% overlap between segments, default number of discrete Fourier transform points) to the detrended fluorescence traces during specified behavioral periods. Statistical significance was assessed at each frequency point using the two-tailed non-parametric Wilcoxon rank-sum test with a significance level of 0.05. For significance, i.e., the PSD of the subthreshold dynamics in one behavioral state (e.g., running) significantly differed from that of another state (e.g., resting), the following notations were used: *p < 0.05, **p < 0.01, and ***p < 0.001 (**Fig. 4j**).

*Characterizations of intercellular cross-correlation, correlation, and coherence.* To determine the similarity and time delay between simultaneously recorded neuron pairs in a given behavioral state, cross-correlation were performed on the same selected periods of filtered fluorescence traces between neuron pairs (100 Hz 5th-order Butterworth low-pass filter with zero-phase shift applied to the detrended traces) (**Fig. 4d**). To estimate the degree of functional connectivity across all neurons in the same field of view, pairwise linear correlation coefficients were computed between each pair of filtered fluorescence traces (using the same preprocessing) during a given behavioral state. The correlation matrices were clustered hierarchically using the “linkage” function (average method) to reveal the heterogeneity in neural activity (**Fig. 4h and Supplementary Fig. 12**). To evaluate the coherence of subthreshold activity between pairs of neurons, subthreshold oscillations within different frequency bands were first extracted from the raw fluorescence trace using a 3rd-order Butterworth bandpass filter. Next, the “mscohere” function (1 s Hamming window, 800 ms overlap between segments, 1024 discrete Fourier transform points) was used to compute the magnitude-squared coherence estimates between all neuron pairs in the same field of view. The mean value of coherence estimates over a specified frequency band was taken as the indicator of coherence strength.

*Phase-locking analysis.* The phase relationship between the spiking and theta-band activity of individual neurons was examined using the Hilbert transform. The “hilbert” and “angle” functions were used to obtain instantaneous theta phases from the extracted theta rhythms, which were then assigned to the timing of detected spikes. A histogram of the phase distribution of spikes relative to the theta cycle was created with a bin size of 0.1 radians (**Fig. 4f and Supplementary Fig.11**). The mean resultant length (MRL) was calculated to reflect the strength of spike phase-locking to theta rhythms using established circular statistics methods [26]. A circular histogram was created with a bin size of 18°, where the length of the red arrow corresponds to phase-locking strength, i.e., the MRL value (**Fig. 4f**).

### Data processing

Images were visualized and processed using Fiji [27]. Unless stated otherwise, all images presented here were unprocessed raw images without smoothing, denoising, and deconvolution.

## CODE AVAILABILITY

MATLAB codes used to generate the results for this study are available from the corresponding author upon reasonable request.

## DATA AVAILABILITY

The raw image sequences are not available at a public repository due to their large size but are available from the corresponding author upon reasonable request.

## ACKNOWLEDGEMENTS

This work was supported by the Beijing Municipal Science & Technology Commission (Z220009), the CAMS Innovation Fund for Medical Sciences (2024-I2M-3-024), and the startup funds from CIBR. The authors thank the CIBR LARC staff for animal care and the CIBR Imaging Core and Instrumentation Core for their support.

## CONTRIBUTIONS

J. W. conceived and supervised the project; J. W. and Z. L. designed the FRAME module; Z. L. and M. H. constructed the microscope, collected, and analyzed the data; Y. C. prepared animals. N. P. D. provided technical support on ScanImage; Z. L., M. H. and J. W. wrote the manuscript with input from all authors.

## SUPPLEMENTARY MATERIALS

**Supplementary Table 1:** Comparison between FACED and FRAME 2PFM for voltage imaging.

**Supplementary Table 2:** Experimental parameters for all imaging datasets.

**Supplementary Table 3:** FRAME-2PFM components list.

**Supplementary Fig. 1.**
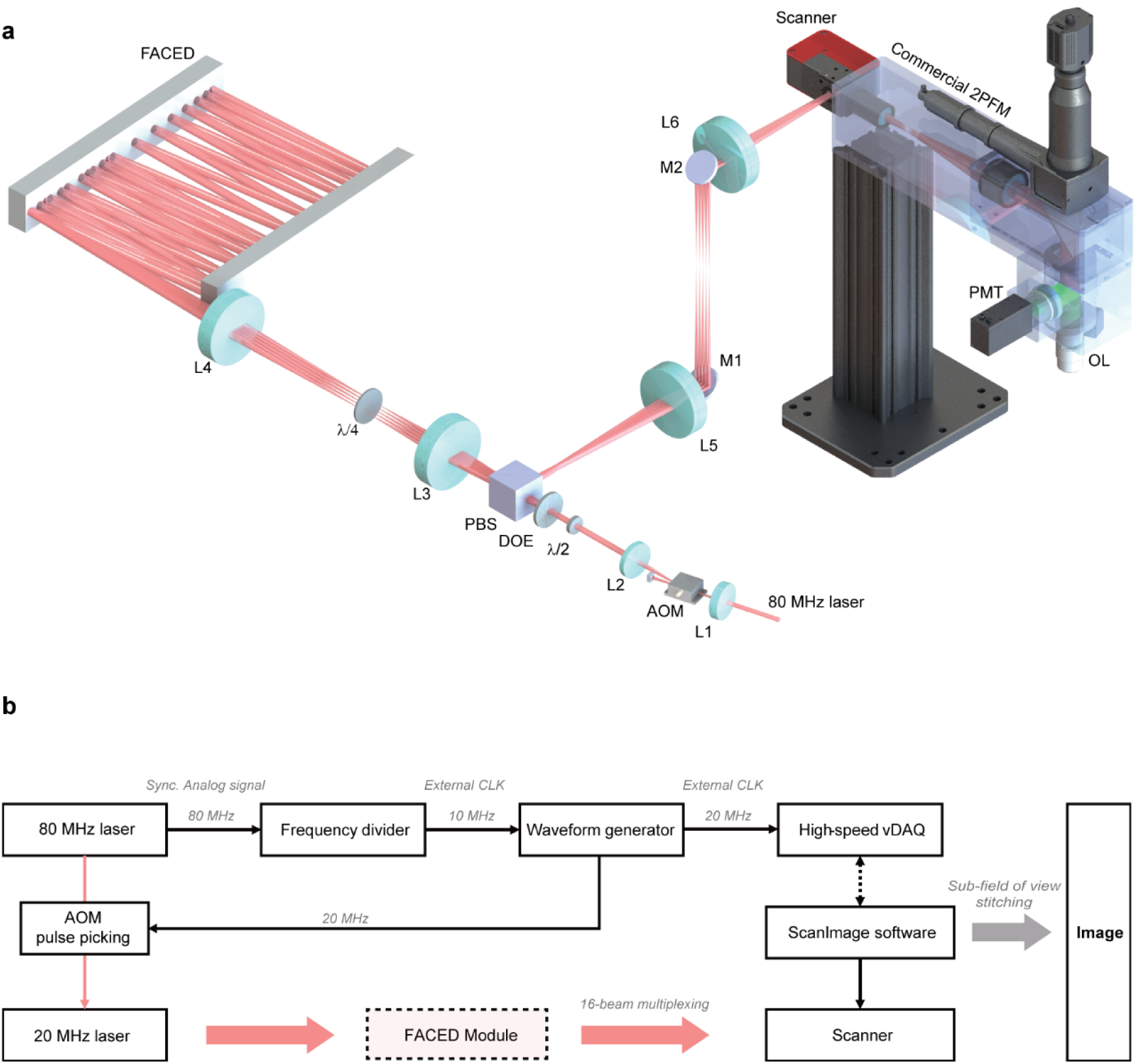
Optical layout and synchronization of FRAME-2PFM. **a**, Optical layout. L1: f = 50 mm; L2: f = 150 mm; L3: f = 250 mm; L4: f = 400 mm; L5: f = 300 mm; L6: f = 250 mm; AOM, acousto-optic modulator; λ/2, Half-wave plate; DOE, diffractive optical element; PBS, polarizing beam splitter; λ/4, quarter-wave plate; M1/M2, mirror; scanner: 12 kHz resonant scanner (x-axis), galvanometer (y-axis); commercial 2PFM: Bergamo II, Thorlabs; OL, objective lens; PMT, photomultiplier tub. **b**, Synchronization control diagram of FRAME-2PFM.

**Supplementary Fig. 2.**
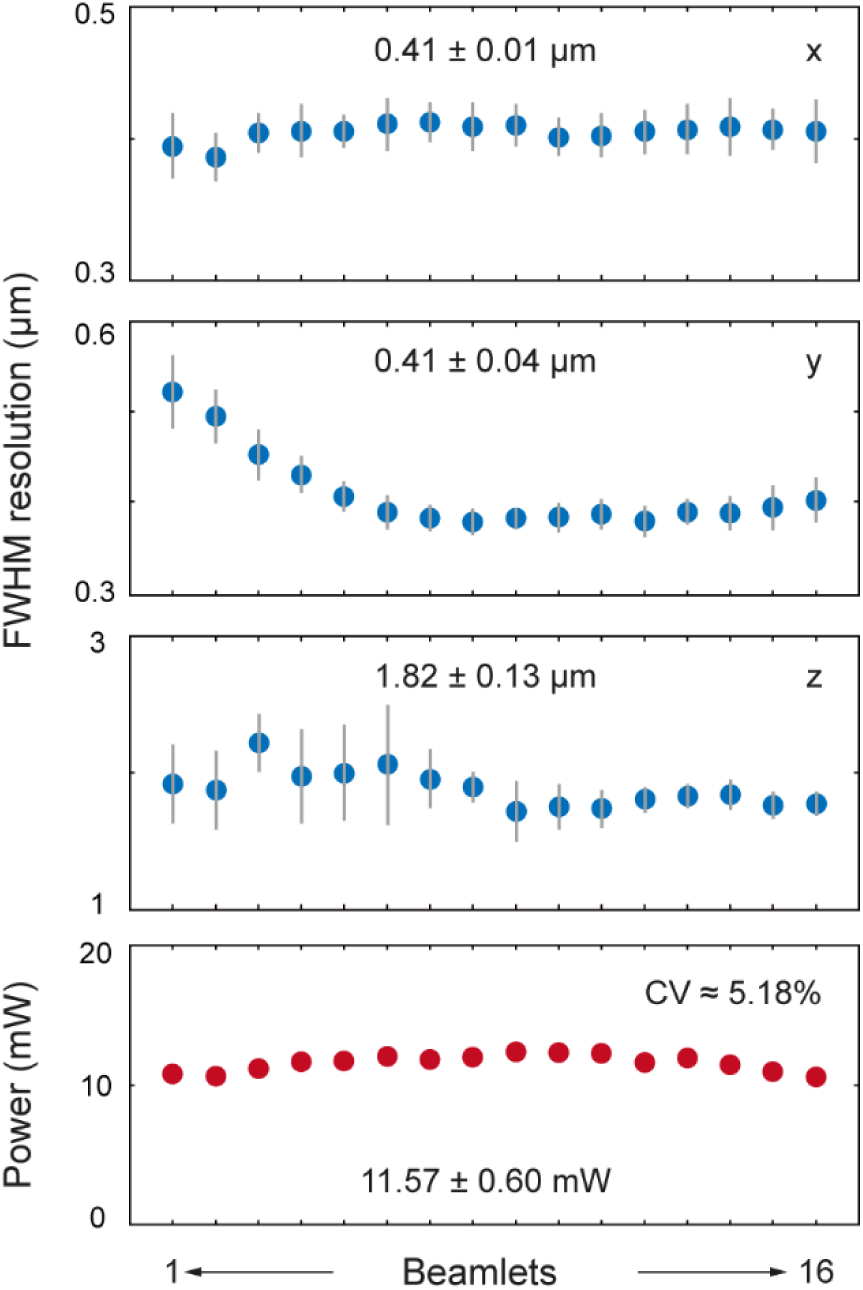
Optical resolution and power uniformity in FRAME-2PFM. Top, optical resolution of FRAME-2PFM (mean ± s.d., n > 10 beads for each sub-field of view); bottom, post-objective power across beamlets.

**Supplementary Fig. 3.**
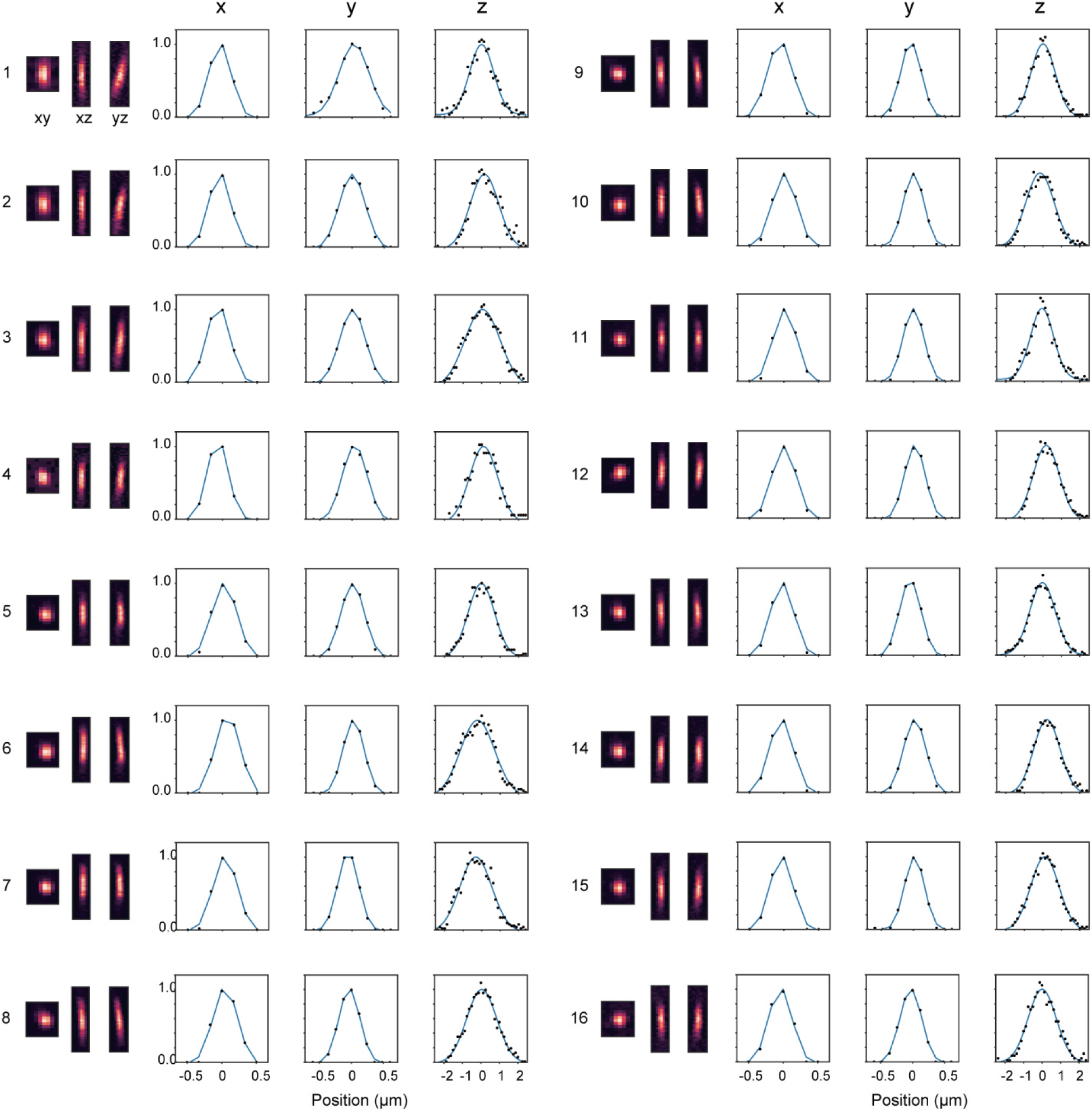
Representative images of 200-nm fluorescent beads in FRAME-2PFM. xy, xz, yz projection views of the beads and PSFs along x, y, and z axes for the 16 sub-fields of view. Gaussian fits are plotted in z-PSFs.

**Supplementary Fig. 4.**
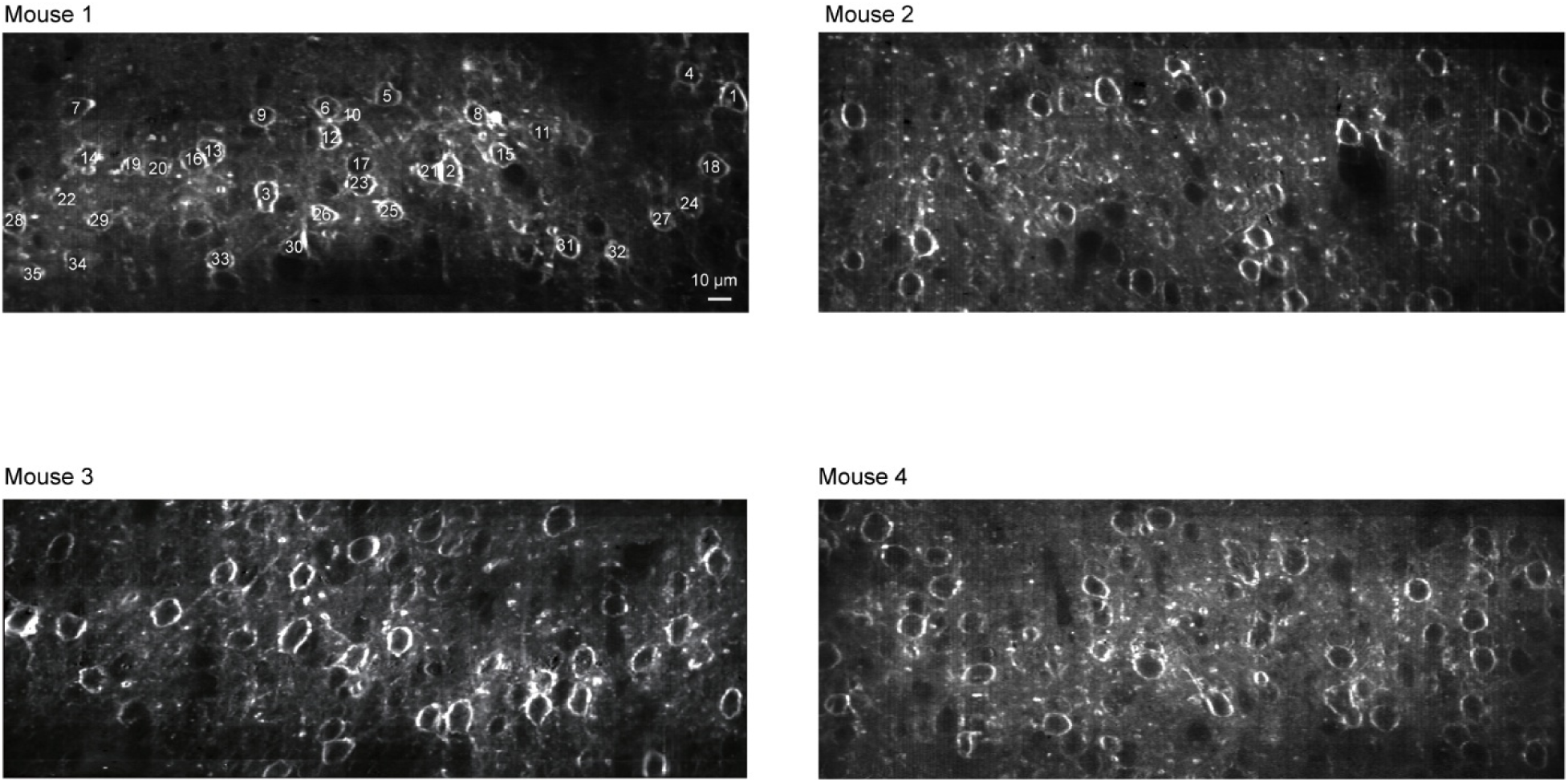
Representative images in V1 with FRAME-2PFM. In the 320 μm × 120 μm field of view, 38 ± 6 neurons were imaged (mean ± s.d., n = 4 mice). Data presented in Fig. 2b were recorded from Mouse 1; the neurons are numbered, and their voltage traces are listed in **Supplementary** Fig. 7.

**Supplementary Fig. 5.**
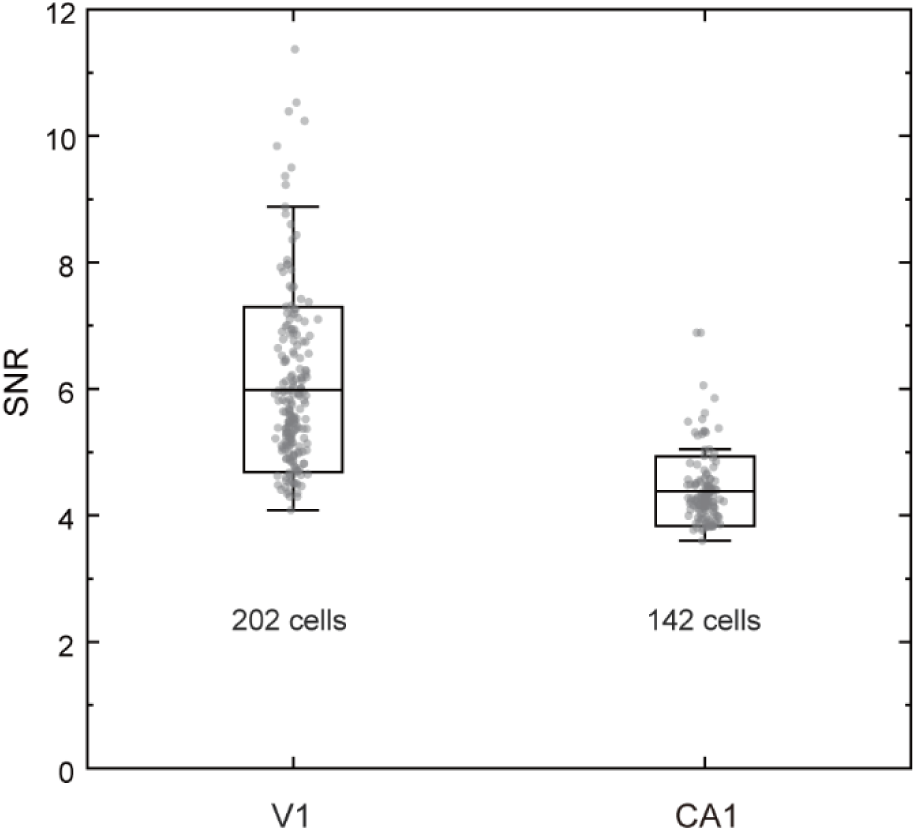
SNR in FRAME-2PFM. SNR in V1 is 6.0 ± 1.3 (mean ± std., n = 202 cells from Fig. 2g); SNR in CA1 is 4.4 ± 0.5 (mean ± s.d., n = 142 cells from **Fig.4b**). Frame rate, 805 fps; field of view, 320 μm × 120 μm; post objective power, 180 mW.

**Supplementary Fig. 6.**
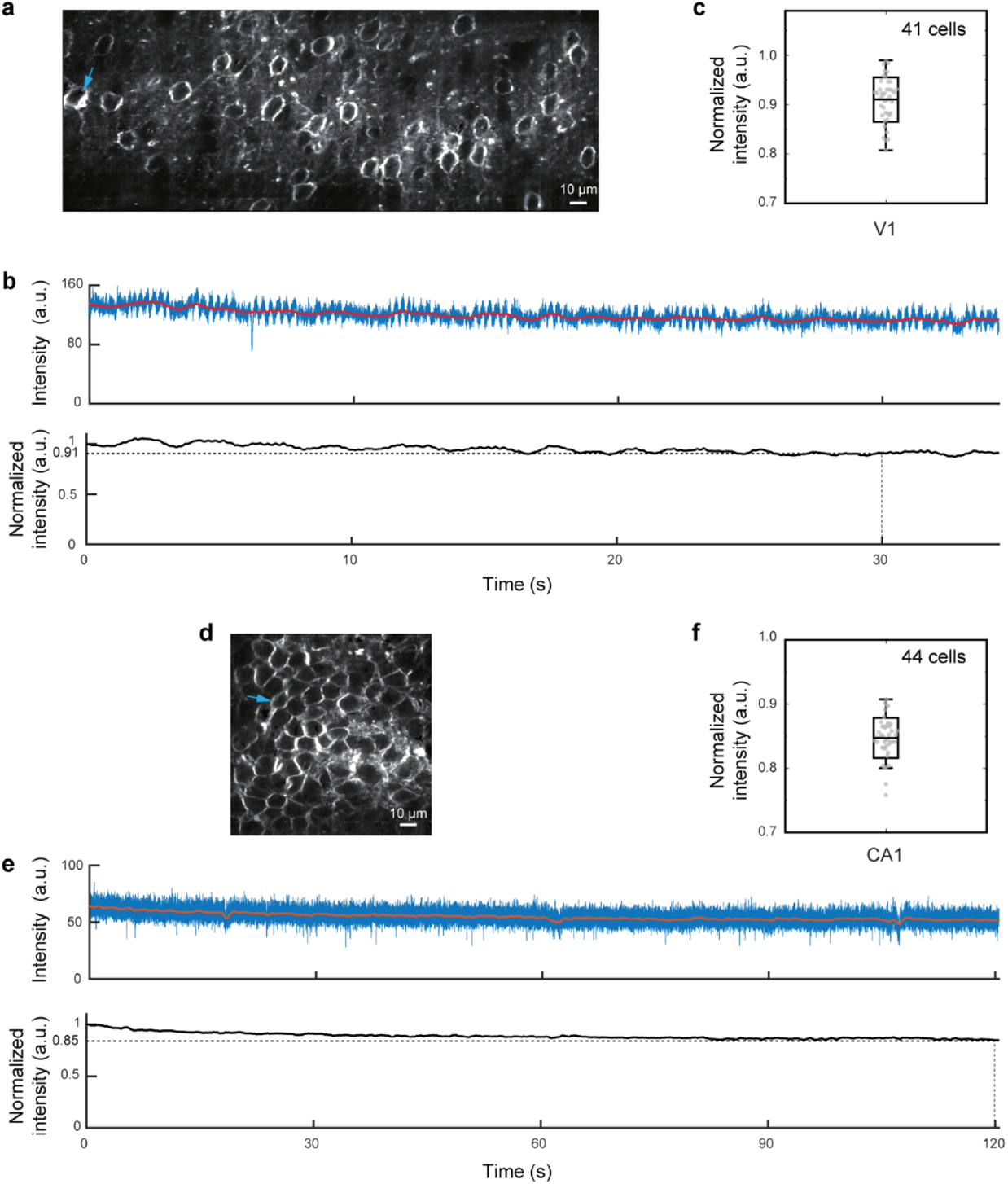
Photobleaching of JEDI-2P-Kv imaged at 805 fps in V1 and CA1. **a**, Imaging parameters in V1: 138 μm imaging depth, 320 μm × 120 μm field of view and 180 mW post objective power. **b**, Top, fluorescence trace of arrow pointed neuron in (**a**). Bottom, normalized mean fluorescence trace of 41 neurons. **c**, The fluorescence intensity in V1 measured after 30 s of continuous imaging was 91.0 ± 0.04% of its initial intensity (mean ± s.d., n = 41cells). **d**, Imaging parameters in CA1: 100 μm imaging depth, 120 μm × 120 μm field of view and 180 mW post objective power. **e**, Top, Fluorescence trace of arrow pointed neuron in (**d**). Bottom, normalized mean fluorescence trace of 44 neurons. **f**, The fluorescence intensity in CA1 measured after 120 s of continuous imaging was 84.0 ± 0.03% of its initial intensity (mean ± s.d., n = 44cells).

**Supplementary Fig. 7.**
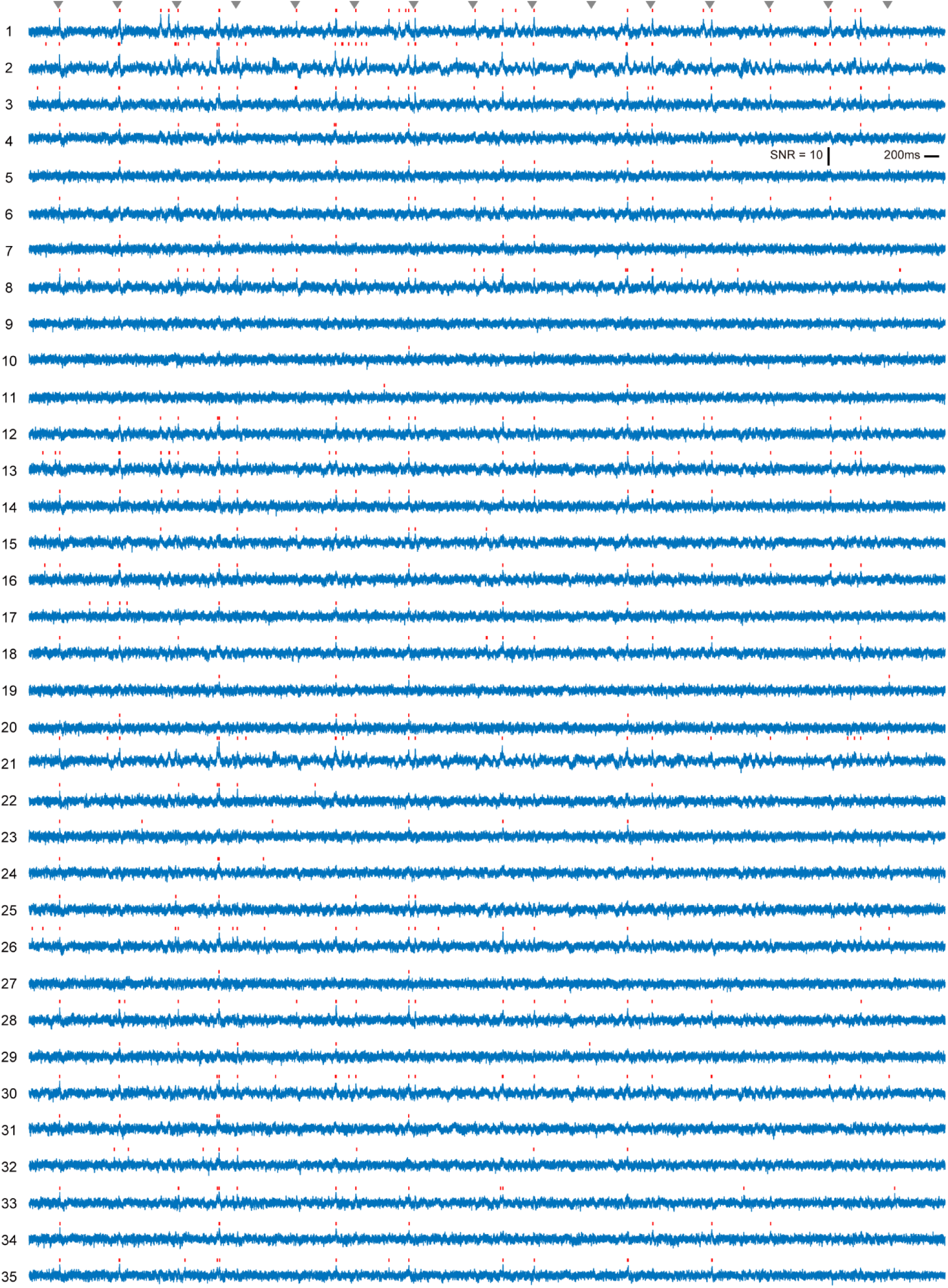
**Voltage traces from 35 neurons recorded in V1 (**Fig. 2b). Gray arrows indicate stimuli; red ticks indicate spikes. Imaging parameters: frame rate, 805 fps; field of view, 320 μm ×120 μm; imaging depth, 120 μm; post-objective power, 180 mW. Light stimulus: 10-ms flash, 2-s interval, 15 flashes per trial. Data presented are from 1 of 9 trials in one animal.

**Supplementary Fig. 8.**
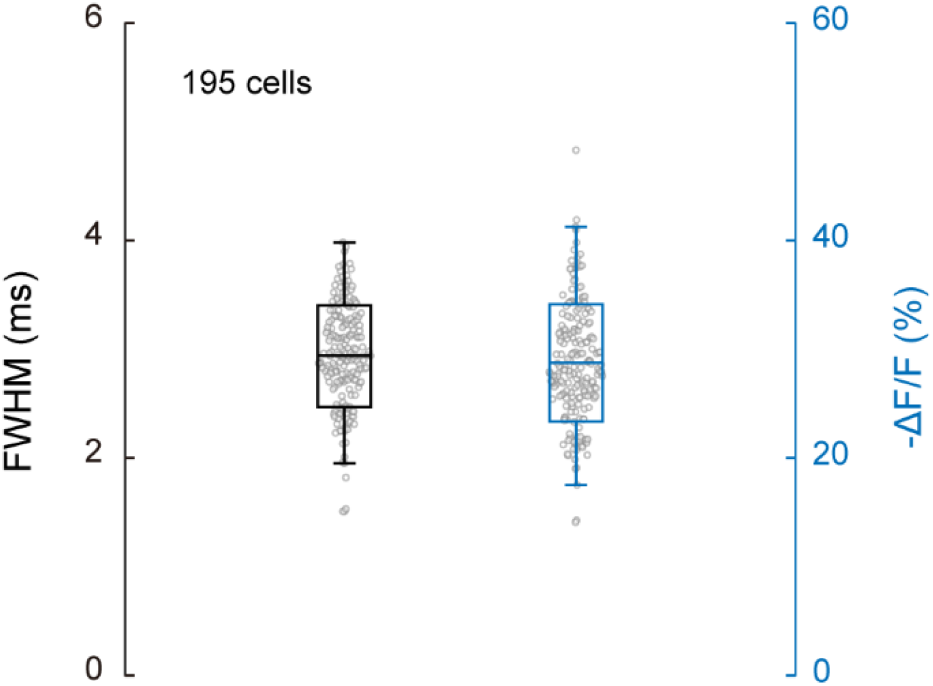
Characteristics of the optical spikes. FWHM and -ΔF/F of optical spikes in V1 (FWHM = 2.93 ± 0.46 ms; ΔF/F = 0.28 ± 0.05; n = 195 cells, mean ± s.d.)

**Supplementary Fig. 9.**
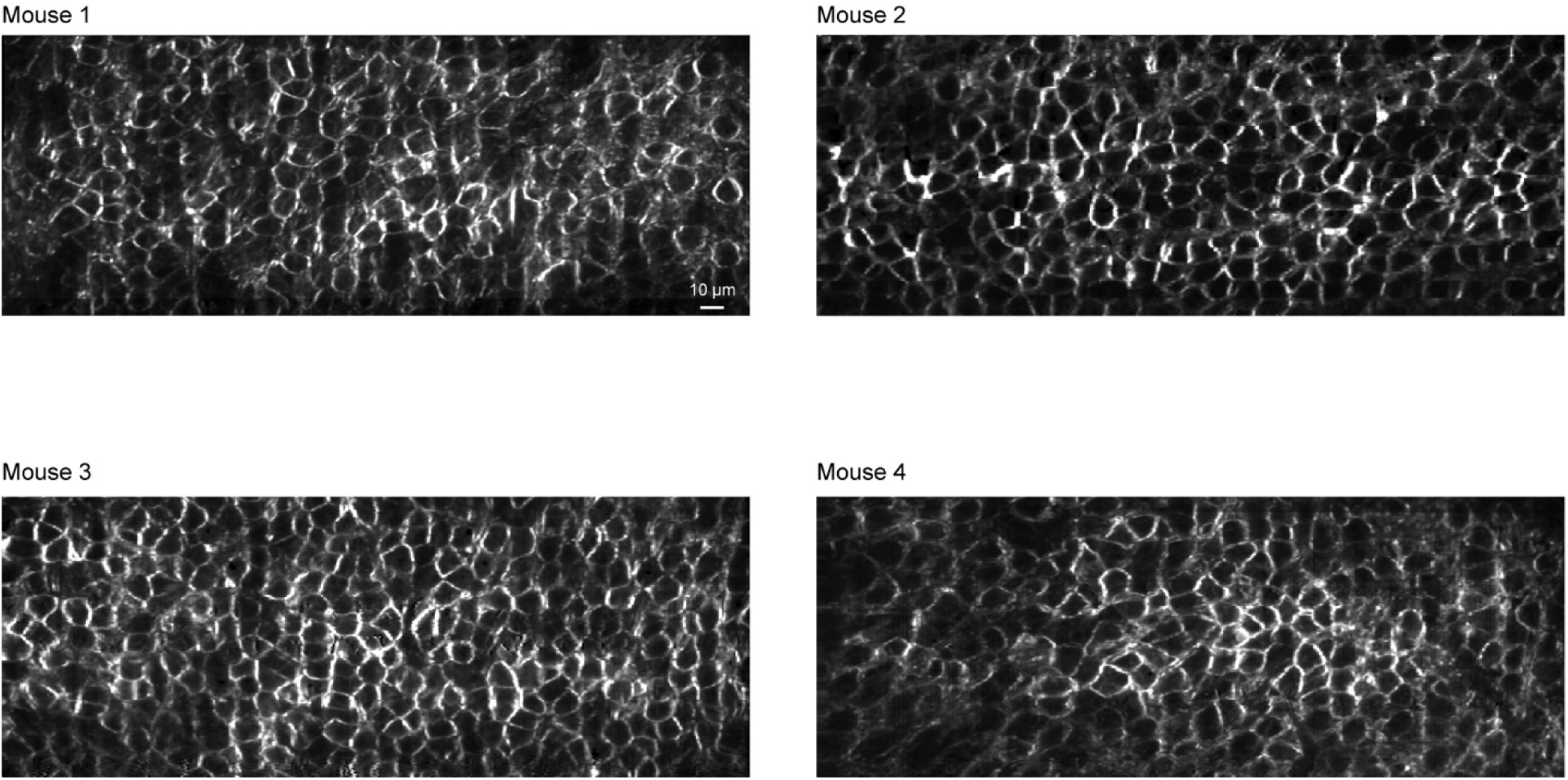
Representative images in CA1 with FRAME-2PFM. In the 320 μm × 120 μm field of view, 203 ± 48 neurons were imaged (mean ± s.d., n = 4 mice). Data presented in Fig. 4 were recorded from Mouse 1.

**Supplementary Fig. 10.**
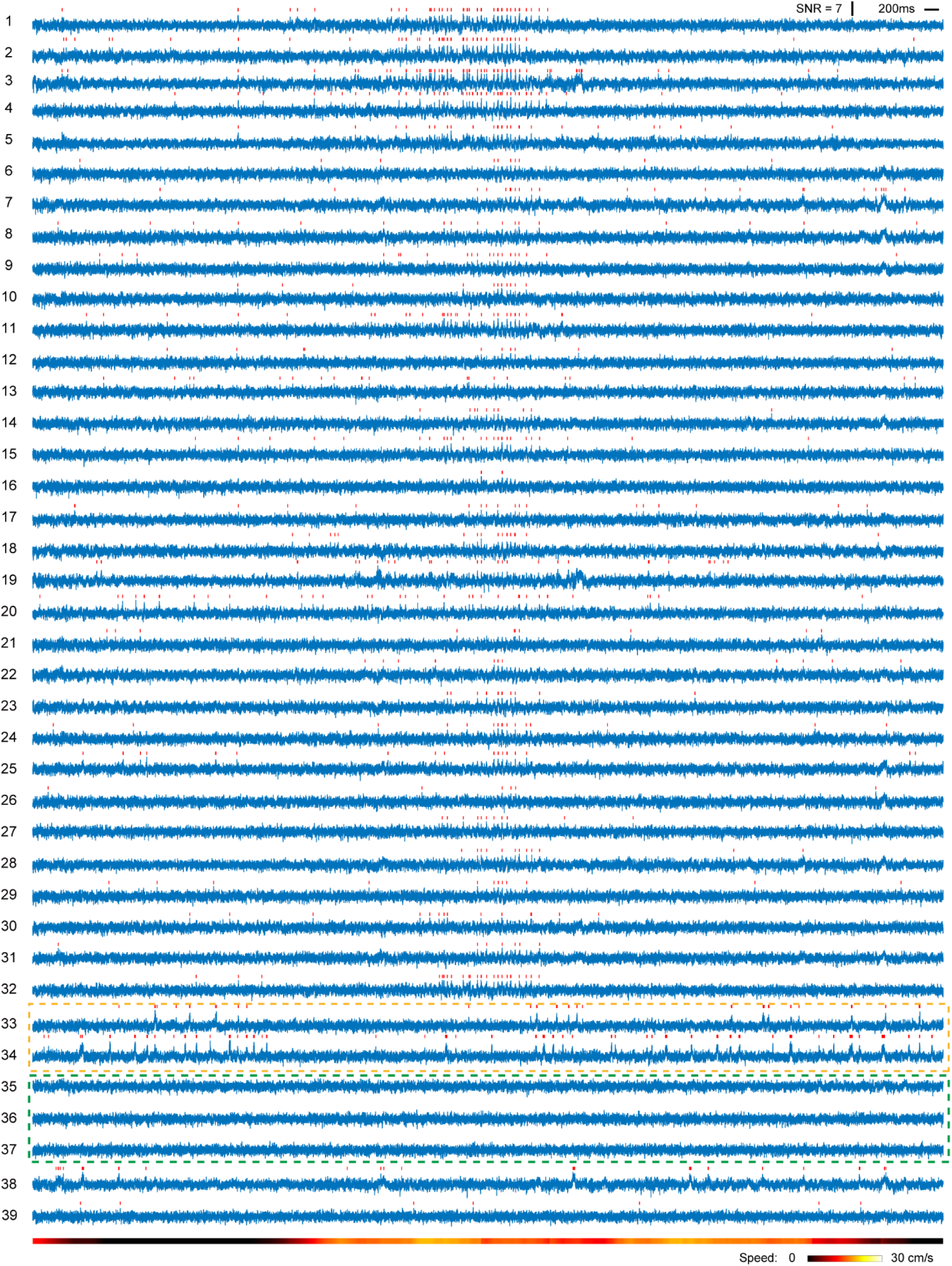
**Voltage traces from neurons recorded in CA1 (**Fig. 4b). Red ticks indicate spikes. Imaging parameters: frame rate, 805 fps; field of view, 320 μm × 120 μm; imaging depth, 100 μm; post-objective power,180 mW. 1–32 are traces of the 32 locomotion responsive neurons; 33–34 and 35–37 are examples traces of neurons showing motion-unrelated spiking activity and no spiking activity; 38–39 are traces from neurons listed in Fig. 4j. Data presented are from 1 of 6 trials in one animal.

**Supplementary Fig. 11.**
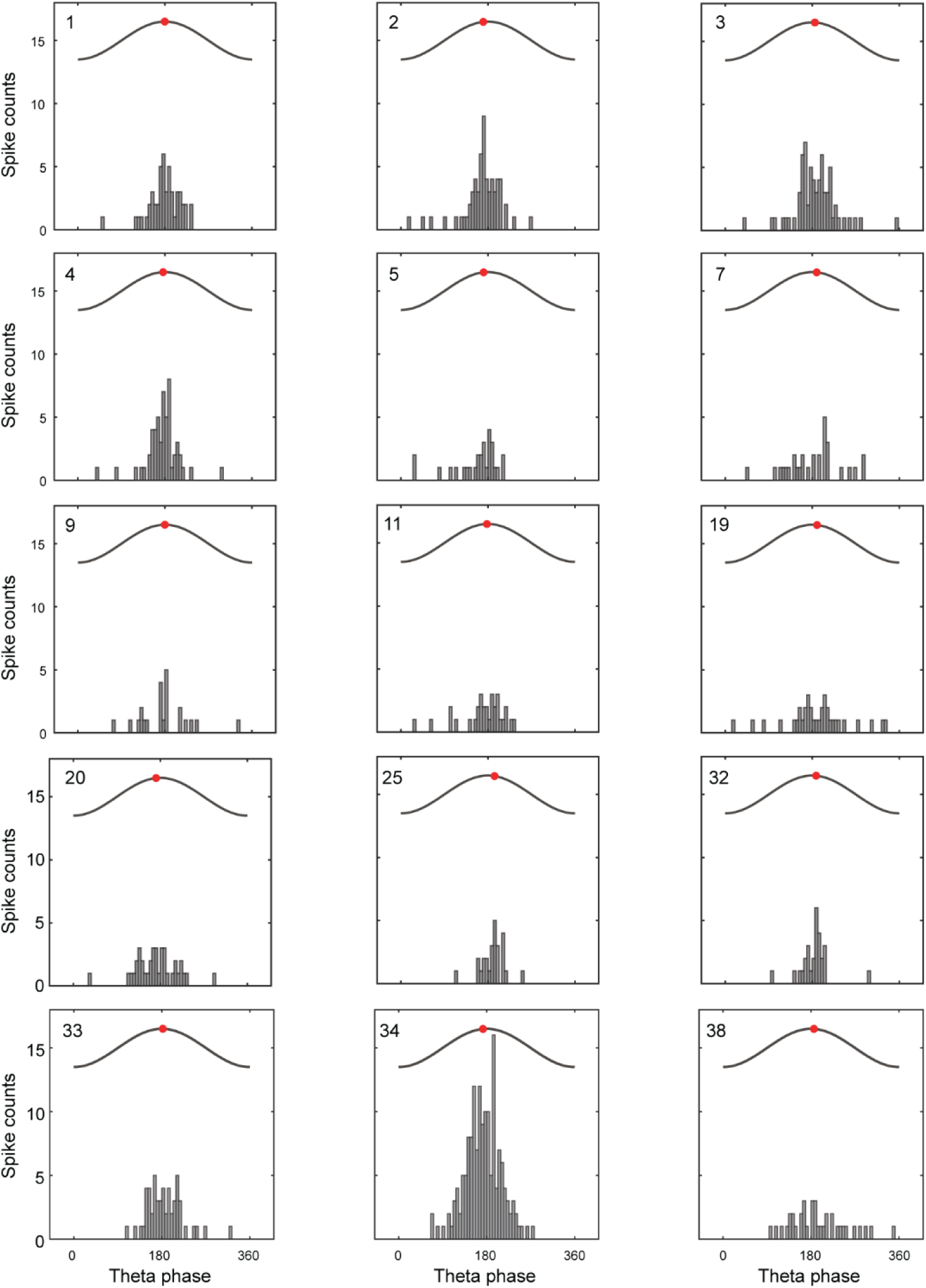
Phase-locking of spiking activity to theta oscillations. Histogram of the phase probability distribution of spikes relative to theta cycles (gray curve) (n = 15 neurons from Fig. 4b). Red dots indicate median phase of the distribution. Cell #: 1, 2, 3, 4, 5, 7, 9, 11, 19, 20, 25, 32, 33, 34, 38.

**Supplementary Fig. 12.**
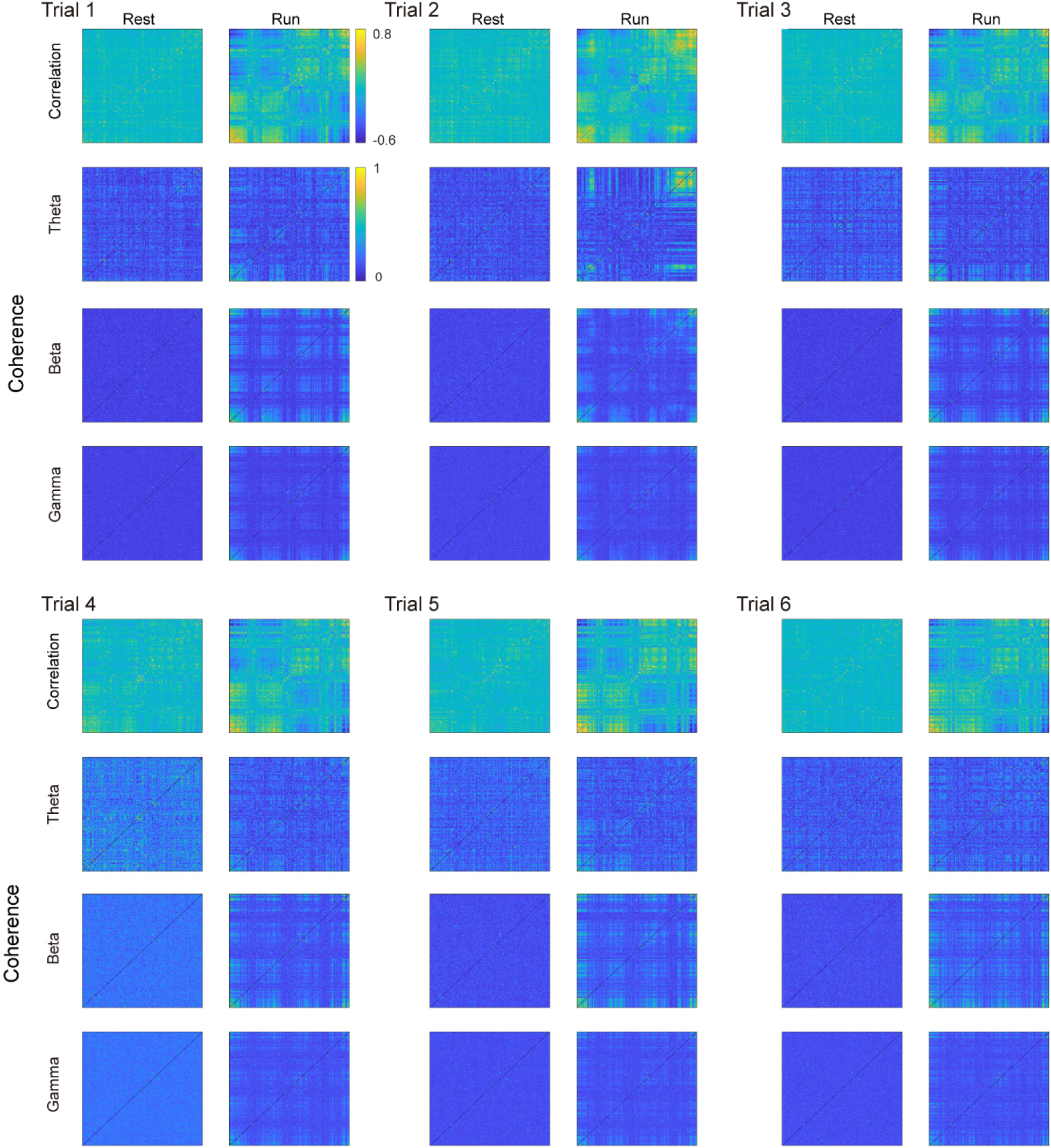
CA1 population activity in different brain states. Correlation and coherence maps of 150 neurons in running and resting states in a single animal. Neurons were ordered based on hierarchical clustering of the running-state correlation matrix in trial 2, and this order was applied to all maps. Only epochs exceeding a 3-s minimum duration were analyzed. Data from the second of six trials are shown in Fig. 4.

**Supplementary Fig. 13.**
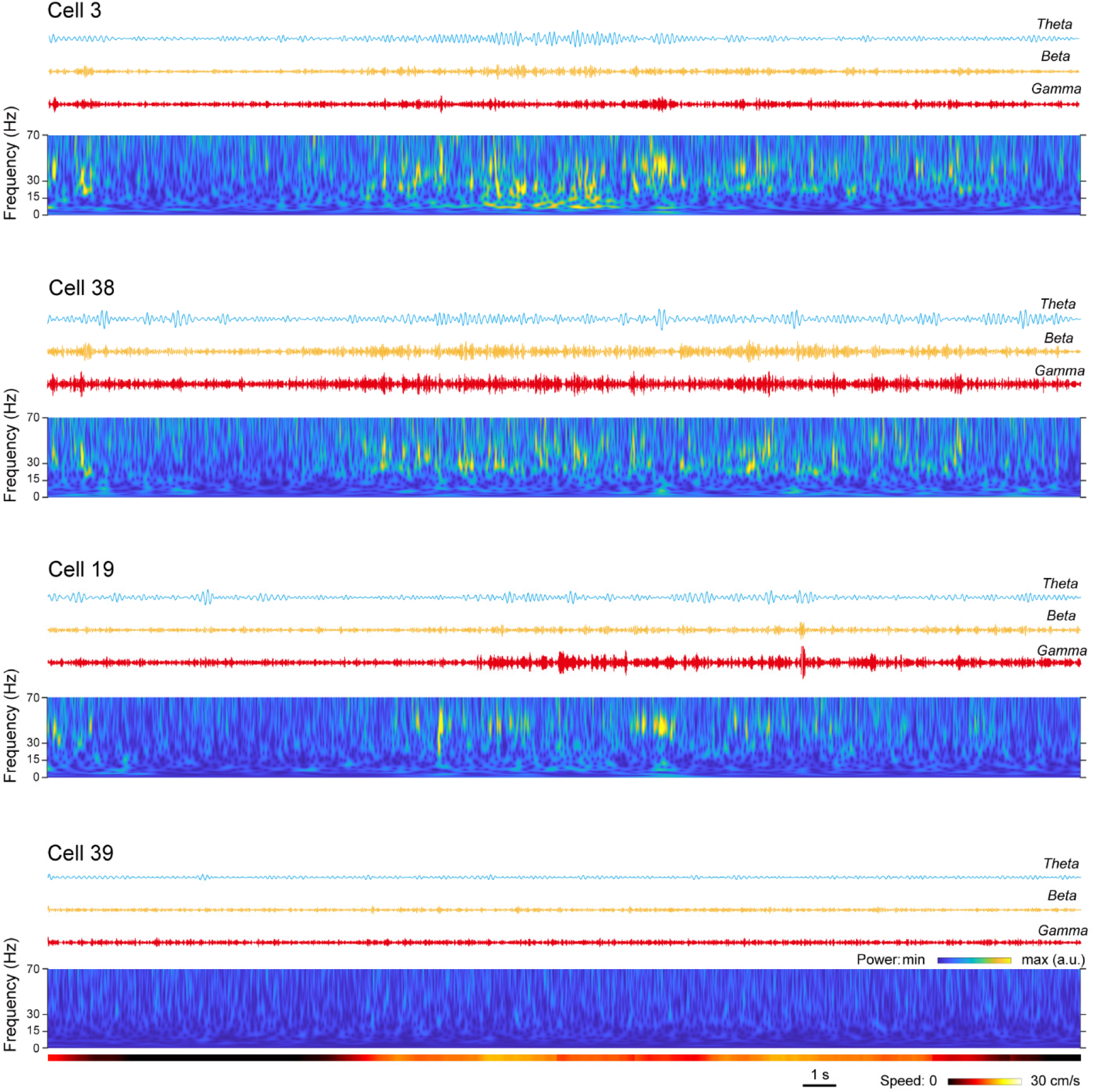
Oscillations and frequency spectrogram of representative neurons. Top traces: theta-, beta-, and gamma-band oscillations extract from the voltage traces of four example neurons in Fig. 4j. Bottom: frequency spectrogram (0–70 Hz).

**Supplementary Fig. 14.**
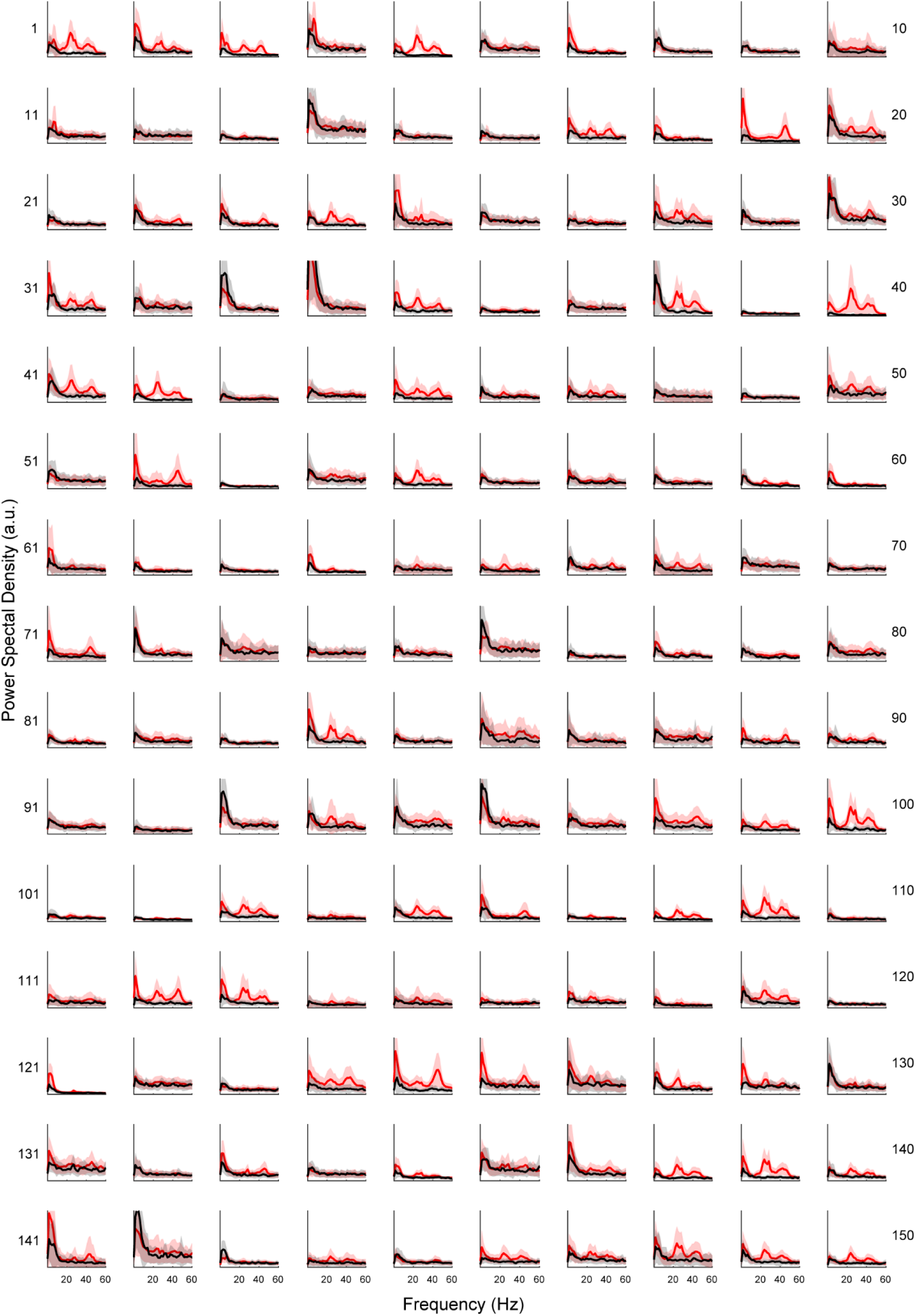
Power spectrums of 150 neurons in. Fig. 4b. Power spectral density was computed from voltage traces recorded during running (red) and resting (black) states (mean ± s.d., n = 6 trials).

**Supplementary Fig. 15.**
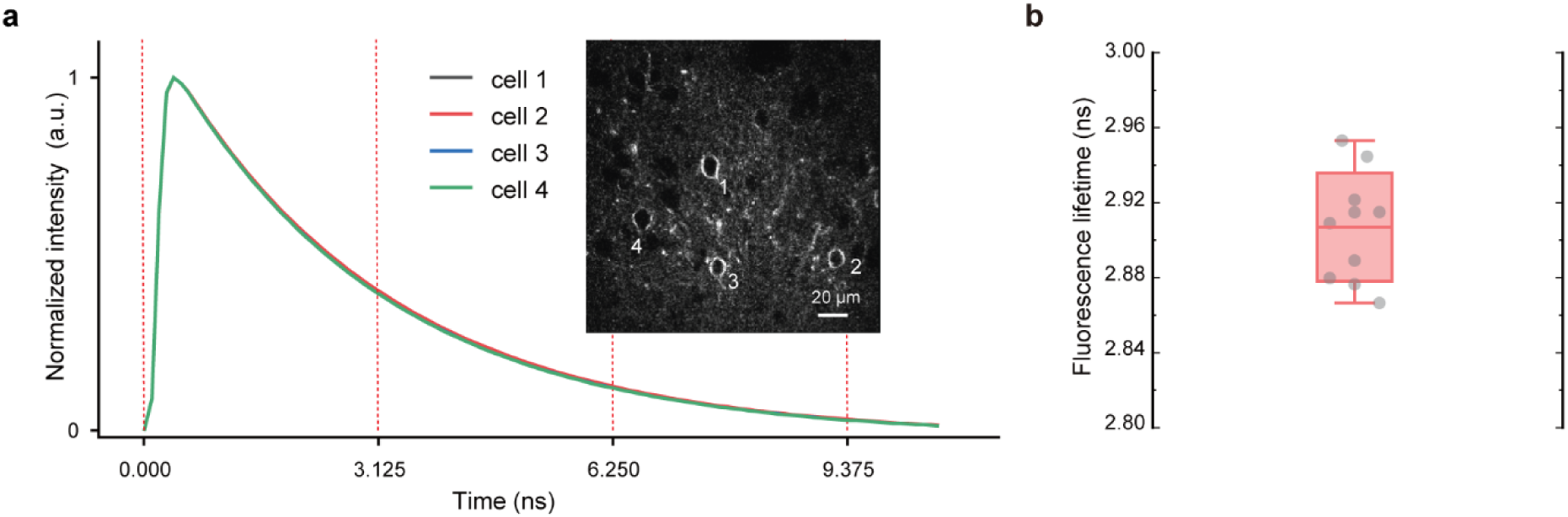
Fluorescence lifetime of JEDI-2P-Kv in vivo. Fluorescence lifetime of JEDI-2P-Kv were measured in vivo in V1 in head-fixed mice using a two-photon fluorescence lifetime imaging microscope (Stellaris 8 Dive/Falcon, Leica). **a**, Image of four neurons and their corresponding fluorescence decay. **b**, Fluorescence lifetime is measured to be 2.91 ± 0.03 ns (mean ± s.d., n = 10 cells).

**Table 1.** Comparison between FACED and FRAME 2PFM for voltage imaging.

| **FACED-2PFM** | **FRAME-2PFM** |
| --- | --- |
| **● Number of pixels along y-axis** |  |
| 80-100 | 384 |
| **● Number of pixels along x-axis** |  |
| 1000 | ~833 |
| **● Imaging throughput (pixel rate)** |  |
| 80–100 MHz | 320 MHz |
| **● Line scan rate** |  |
| 1 MHz | 384 kHz |
| **● Laser source** |  |
| 1 MHz / microjoule pulse energy | 80 MHz/ nanojoule pulse energy |
| Notes: the number of pixels and pixel rate are calculated without accounting for scan dead time at 1000 fps; the pixel rate represents the maximum achievable rate under the condition of one pulse per pixel; the FACED scanner is configured to scan y-axis for comparison purposes; the 80 MHz laser in FRAME-2FPM is reduced to 20 MHz for multiplexing. |  |

**Table 2.** Experimental parameters for all FRAME-2PFM in vivo imaging datasets.

| Brain area | Figure # | Post-objective power (mW) | Imaging time (s) | Field of view ($\mu\text{m}^2$) | Frame rate (fps) | Imaging depth ($\mu\text{m}$) | Number of cells |
| --- | --- | --- | --- | --- | --- | --- | --- |
| V1 | Fig. 2b<br>SI Fig. 4 (mouse 1) | 180 | ~31 | 320 x 120 | 805 | 120 | 35 |
|  | Fig. 3a | 180 | ~30 | 96 x 120 | 1006 | 60 | 1 |
|  | SI Fig. 6a<br>SI Fig. 4 (mouse3) | 180 | ~37 | 320 x 120 | 805 | 138 | 41 |
|  | SI Fig. 6d | 180 | ~120 | 120 x 120 | 805 | 100 | 44 |
|  | SI Fig. 4 (mouse 2) | 180 | ~31 | 320 x 120 | 805 | 150 | 31 |
|  | SI Fig. 4 (mouse 4) | 180 | ~31 | 320 x 120 | 805 | 100 | 44 |
| CA1 | Fig. 4b<br>SI Fig. 9 (mouse 1) | 180 | ~31 | 320 x 120 | 805 | 100 | 150 |
|  | SI Fig. 9 (mouse 2) | 140 | ~30 | 320 x 120 | 1006 | 70 | 222 |
|  | SI Fig. 9 (mouse 3) | 180 | ~32 | 320 x 120 | 928 | 100 | 260 |
|  | SI Fig. 9 (mouse 4) | 180 | ~31 | 320 x 120 | 805 | 120 | 180 |

Note: all FRAME-2PFM images presented in this work are mean intensity projections from their respective imaging stacks.

**Table 3.**
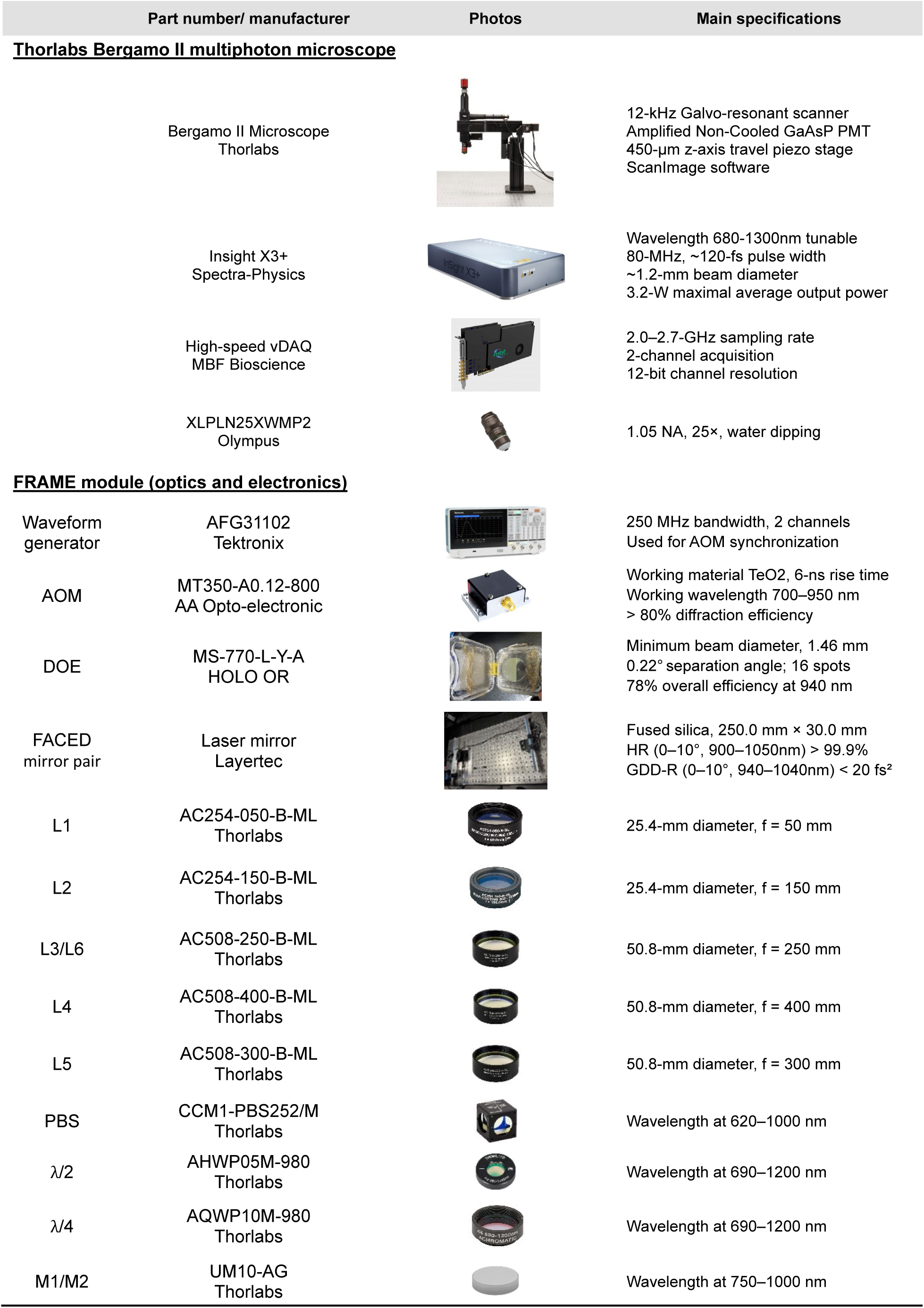
FRAME 2PFM components list.

